# Binding affinities for 2D protein dimerization benefit from enthalpic stabilization

**DOI:** 10.1101/2025.01.16.633485

**Authors:** Adip Jhaveri, Smriti Chhibber, Nandan Kulkarni, Margaret E Johnson

## Abstract

Dimerization underpins all macromolecular assembly processes both on and off the membrane. While the strength of dimerization, K_D_, is commonly quantified in solution (3D), many proteins like the soluble BAR domain-containing proteins also reversibly dimerize while bound to a membrane surface (2D). The ratio of dissociation constants, 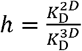, defines a lengthscale that is essential for determining whether dimerization is more favorable in solution or on the membrane surface, particularly for these proteins that reversibly transition between 3D and 2D. While purely entropic rigid-body estimates of *h* apply well to transmembrane adhesion proteins, we show here using Molecular Dynamics simulations that even moderate flexibility in BAR domains dramatically alters the free energy landscape from 3D to 2D, driving enhanced selectivity and stability of the native dimer in 2D. By simulating BAR homodimerization in three distinct environments, 1) solution (3D), 2) bound to a lipid bilayer (2D), and 3) fully solvated but restrained to a pseudo membrane (2D), we show that both 2D environments induce backbone configurations that better match the crystal structure and produce more enthalpically favorable dimer states, violating the rigid-body estimates to drive *h*≪*h*_*RIGID*_. Remarkably, contact with an explicit lipid bilayer is not necessary to drive these changes, as the solvated pseudo membrane induces this same result. We show this outcome depends on the stability of the protein interaction, as a parameterization that produces exceptionally stable binding in 3D does not induce systematic improvements on the membrane. With *h* lengthscales calculated here that are well below a physiological volume-to-surface-area lengthscale, assembly will be dramatically enhanced on the membrane, which aligns with BAR domain function as membrane remodelers. Our approach provides simple metrics to move beyond rigid-body estimates of 2D affinities and assess whether conformational flexibility selects for enhanced stability on membranes.

## I. Introduction

The formation of reversible protein dimers from monomeric building blocks during membrane trafficking ^1 2, 3^, signalling^4-6^, and pattern formation^7, 8^ is often of only moderate to weak stability, ranging from *K*_*D*_^′^*s* in the high nanomolar^9, 10^ to high micromolar^11, 12^, respectively. Forming a large population of dimers is then improbable without high concentrations that reach the micromolar range, were these proteins in isolation. However, many of these proteins also reversibly localize to membranes^13^, often using domains such as the BAR domain studied here that directly bind membrane lipids^14-16^. These so-called peripheral membrane proteins will continue to diffuse on the membrane. Any two such localized monomers that could bind in 3D can now undergo reversible dimerization in the effectively 2D environment on the membrane. How will this surface localization alter the strength and thus probability of dimerization? The binding free energy of this reversible dimerization, Δ*G*^2*D*^ must change compared to Δ*G*^3*D*^ due to entropy loss. At a minimum, this includes the loss of a search dimension and the restriction of monomer and dimer rotational freedom upon surface adhesion. A particularly useful metric for comparing these changes is the lengthscale 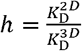, as these dissociation constants are independent of system size, whereas Δ*G* is not (the standard state in 3D is different than 2D). These metrics are related via 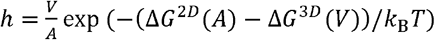, where *V* and *A* are the volume and surface area domains for measuring Δ*G, k*_B_ is Boltzmann’s constant and *T* is temperature. How large do we expect this lengthscale range to vary due to additional entropic changes to backbone or sidechain sampling, protein-membrane interactions, or enthalpic changes? Here we generalize a statistical mechanical theory ^17,18^ of the *h* lengthscale to consider non-rigid proteins, with applications to peripheral membrane proteins. We specifically show using MD simulations^19^ coupled to enhanced sampling^20, 21^ that additional entropic and enthalpic factors beyond the rigid-body approximation can widely tune this lengthscale for the BAR-domain protein LSP1, which is known to dimerize in both 3D and 2D^22-24^.

The magnitude of the lengthscale *h* is especially critical for peripheral membrane proteins because they reside in the cytoplasm but can reversibly transition to membranes. This lengthscale is thus a key determinant in whether dimers will form more favorably in the proteins cytoplasm (3D) or on the membrane (2D). For comparing a *population* of *N* dimer-forming in a 3D volume *V*_sys_ vs a 2D area *A*_sys_, we are guaranteed to observe more dimers in 2D if 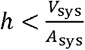, which follows from the definition of the dissociation constant,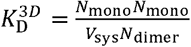 or LeChatelier’s principle^25^. More generally, for a population of proteins that can reversibly transition from *V*_sys_ to *A*_sys_ (via lipid binding, for example), dimerization will be enhanced relative to 3D binding if 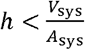, with the extent of enhancement depending in large part on the affinity of proteins for the membrane^26^. For cells of radius 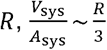, dimerization is enhanced on membrane surfaces when is below the ∼0.5-10 *μm* scale. This dimensional reduction from a 3D to a 2D search for binding partners can thus dramatically enhance self-assembly^26^, in addition to the well-established impact on search timescales^27, 28^, but it depends on the thermodynamics of the specific binding pair.

The magnitude that has been previously measured for is roughly the nanometer scale, thus favoring dimerization on the membrane. However, several protein-specific and environmental factors will impact the order-of-magnitude. Simulation^29^, rigid-body approximations^29^ or experiment^30^ have measured to be on the nanometer scale for two proteins that bridge across two membranes, but the separation distance between the membranes and their roughness can alter the ratio by ∼10 fold, with lower values reaching *h* ∼ 0.1 nm^17, 18^. Peripheral membrane proteins, including the BAR domain, lack direct experimental quantification of 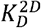 because they frequently transition between 2D and 3D, convoluting the also measure nanometer scales^31-33^, with estimates of *h* =8*nm* -18*nm*^33^. Larger values of binding measurements. However, receptor-ligand interactions restricted to the same membrane ≈30*nm*^34^ were inferred for the clathrin-clathrin dimerization interaction based on a best-fit studies support in the 0.1-50*nm* range for functionally relevant 2D interactions. Physically model to in vitro fluorescence of clathrin assembly on membranes in time^35^. Collectively, these both larger and smaller ratios are possible, as we see here, and the BAR domain we study here represents an excellent test case because of its distinct binding structure. Unlike the systems above, the BAR domain dimer interface is directly adjacent to its membrane binding interface, within the same domain^14, 36^. This proximity to the membrane reduces orientational sampling and likely reduces translational motion in *z*. Based on purely rigid-body approximations, we thus predict (and find) a lower *h* < 0.1*nm*. However, if the rigid-body approximation fails, conformational changes could push *h* in either direction. For example, if membrane binding occludes the protein-protein interface, we expect the ratio to rise to effectively *h →* ∞, since no specific binding will occur in 2D. Alternatively, if membrane binding induces conformational changes in the proteins^37^ that creates or significantly improves a stable binding interface, could fall well below the sub nanometer regime.

To test these effects, we simulated the LSP1 dimerization process in the MARTINI 2.0 and 3.0 forcefields. Simulation and theory have key advantages for providing us the first quantitative estimates of 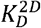 and *h* for this essential family of protein domains. We can isolate and 3.0 forcefields. Simulation and theory have key advantages for providing us the first the 2D dimer and monomer ensembles by readily discarding simulation intervals when the proteins dissociate from the membrane, and we avoid oligomerization by simulating only two proteins. Importantly, the simulations provide structural and energetic details necessary to assess the microscopic origins of *h*, allowing us to quantify contributions to *K*_D_ from diverse BAR dimer bound states, as done in previous solution (3D) simulations^16^. Previous simulation studies of 2D dimerization (bridging membranes) used rigid protein domains with limited interdomain flexibility and no solvent^17, 18, 29^. By using the relatively high-resolution MARTINI coarse-grained force-field, we retain explicit solvent and flexibility of the protein backbone and side chains. This allows us to consider entropy changes arising from far more degrees of freedom and relax the assumption that the enthalpy of the interaction is unchanged. In the MARTINI 3.0 forcefield (LSP_M3.0)^38^, which more accurately predicts binding free energies^38, 39^, we find *h* is extremely low, reaching sub-femtometer levels, indicating much more stable 2D dimerization. In contrast, simulations in MARTINI 2.0 (LSP_M2.2p) measure *h* in the micrometer range, indicating weakened binding in 2D. While this clearly indicates force-field dependencies^40^, the rich data from the simulations allows us to attribute these quantitative and qualitative variations between the two dimers to differential contributions of enthalpic vs entropic terms in both cases. With simulations, we can also control for the impact of explicit protein-lipid interactions and curvature variations on the measured 2D behavior by constructing a 2D pseudo membrane using only geometrical restraints in water. Remarkably, for the LSP_M3.0 dimer, we find that much of the improved stability in protein conformations is induced already with the pseudo membrane and therefore is not dependent on an explicit protein-lipid interface. Our analysis below provides a formal background for quantifying the rigid-body approximation to and exploiting simulations to assess its accuracy in predicting how peripheral membrane proteins select for specific dimers in 2D environments.

## II. THEORY

### II. A Theoretical background on 3D and 2D binding thermodynamics

As essential background, we introduce established thermodynamic equations^41^. The equilibrium constant for a pair of protein monomers binding to one another in solution (3D) is given by:

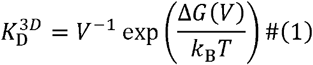

Where *k*_B_ is Boltzmann’s constant, *V* is the system volume, *T* is the temperature, and Δ*G* is the Gibb’s free energy difference between two states *s* ∈ (*bnd, unb*), the bound and unbound monomers, Δ*G* (*V*)= *G*_*bnd*_ (*V*)-G_*unb*_ (*V*) at a constant *NPT, N* particle number and *P* pressure. The Gibb’s free energy for a state *S* depends on the Helmholtz free energy *F*_s_ via *G*_*s*_ = F_*s*_ + *PV*_s_, thus the free energy difference is:

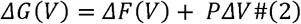

where *ΔF* and *ΔV* report the difference between the bound and unbound states. The contribution of in Eq. 2 is typically negligible compared to the change in internal energy and entropy of the bound and unbound states, so we assume it can be neglected^41^, *V*_*bnd*_ ≅ V_*unb*_ and Δ*G* (*V*) ≅ Δ*F*(*V*). The primary determinant of the binding free energy Δ*G* is through Δ*F*, which is calculated from the canonical partition function *Q*^42^. If we assume the momentum integrals are the same for the bound and unbound ensembles, the ratio of the bound and unbound partition functions *Q* _*bnd*_ /*Q*_*unb*_ will thus reduce to the ratio of the bound and unbound configuration integrals (see SI):

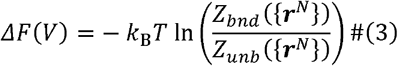

Both *Z*_*bnd*_ and *Z*_*unb*_ depend on the system volume, but most of those factors cancel out when taking their ratio, and{**r**^*N*^} is all atomic coordinates in the system. In 3D, the configurational sampling of two monomers in the unbound state (versus one dimer in the bound state) preserves the dependence of *ΔF* on *V*.

The binding free energy between a pair of protein monomers that are restricted to a membrane/surface no longer scales with the volume of the system, but rather with the surface area *A*. As the *z*-dimension (height) of the system increases, the Δ*G* should not change, as the additional solvent volume is not sampled by either the bound or unbound monomers and thus does not contribute to the relative free energy. Thus, we must assert the system as being restricted to effectively 2D sampling; the proteins cannot unbind from the membrane. The 2D dissociation constant is then given by:

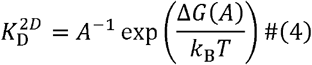

Since the molecules still exist in 3D space, the partition functions of Eq. 3 are still defined over the same protein degrees of freedom as in 3D, but restrictions to the *z*-dimension of the proteins allowed within that volume enforces a 2D search. These restrictions to the surface then further limit rigid-body orientational sampling (the Euler angles: yaw, pitch, and roll), as proteins align a specific interface to the membrane. However, they may also restrict configurational sampling of the protein’s internal degrees of freedom via backbone or side-chain rearrangement, which is not captured by rigid-body degrees of freedom.

The ratio of 3D and 2D equilibrium constants for a pair of protein monomers gives:

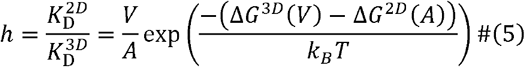

Inserting Eq. 2 and Eq. 3 into this formula, and ignoring/cancelling the *P Δ V* terms for 2D and 3D produces:

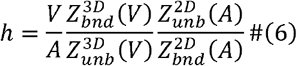

With a series of assumptions about these partition functions, we will arrive at the rigid-body approximation to Eq. 6 that is readily calculated simply from the fluctuations of the monomers restricted to 2D.

### II. B Rigid-body approximation for *h*

To simplify *h* defined in Eq. 6^18, 43^, we define the partition functions in the SI Methods as a function of all coordinate degrees of freedom, where we use similar notation to Ref^41^ and ^44^ (Fig S1). The key difference of the 2D system compared to the 3D system is we now consider both proteins 1 and 2 to be bound to the membrane surface, with the membrane oriented in the x-y plane such that its normal typically points in the z-direction (Figure 1). Unlike in 3D, the six integrals over the translational position ***R***_1_ =x_1_,y_1_, z_1_ and rigid-body orientation Ω= θ_1_,Φ_1_, Ψ_1_ of protein 1 (and 2) in the volume are now restricted by the membrane in heightz_1_, the pitch θ_1_, and the roll Φ_1_ (Fig 1, Fig S1). The other three integrals over *x*_1_, *y* _1_, and the yaw Ψ are unrestrained just as in 3D. With a series of systematic assumptions (see SI for derivation) we separate the potential energy contributions from height, pitch, and roll of 2D bound and unbound monomers from the potential energy contributions of all internal atomic coordinates of monomers and solvent *r*_1_,*r*_*hyd*_, *r*_2_ and from the potential energy due to the translational position and orientation of protein 2 relative to protein 1 (*R*_21_, Ω_2_). We arrive at the approximation:

**Figure 1.**
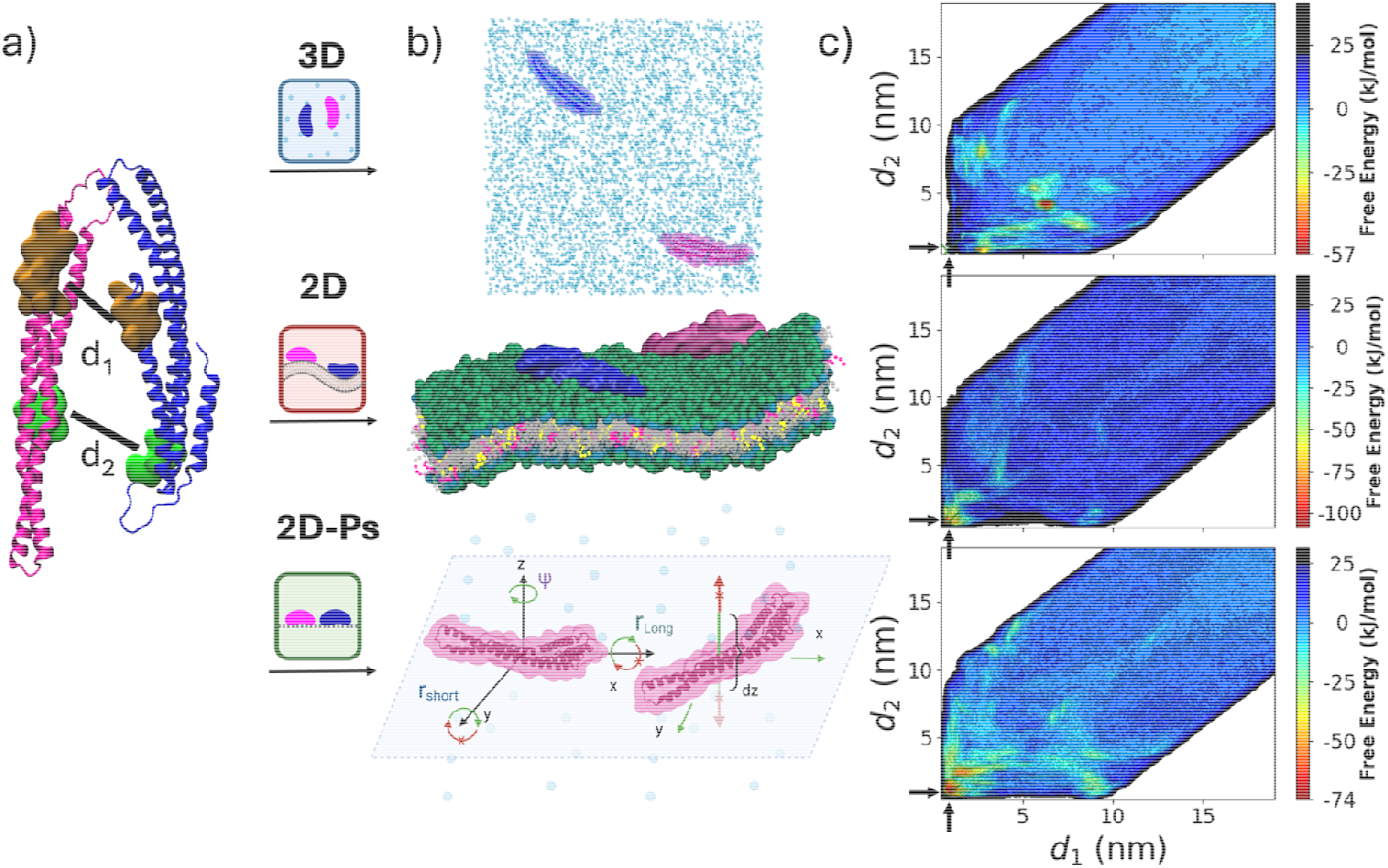
Overview of simulated dimer environments and analysis approach. a) Two LSP1 BAR domain monomers with the metadynamics collective variables d_1_ and d_2_ defined based on the brown and green patch separations. b) Dimerization is simulated in three different environments: solution (3D), membrane bilayer (2D) and a pseudo-membrane 2D plane (2D-Ps) enforced by geometrical restraints that limit the translational motion in, and the orientational sampling around the long and short axis of each monomer. The bilayer has 80% POPC (gray), 10% POPS (pink) and 10% PIP_2_ (yellow). c) For each environment we performed a single long MD simulation with enhanced sampling via metadynamics to build the free energy surface (FES) with respect to the collective variables d_1_ and d_2_. The native/crystal dimer is indicated by the black arrows at d_1_=0.77nm, d_2_=0.86nm. FES is shown for LSP_M3.0 (MARTINI 3.0), and color bars are per environment. We repeated the same approach for LSP_M2.2p (MARTINI 2.0). The binding affinities,, and are obtained by integrating the FES over the bound versus unbound states (see Methods).

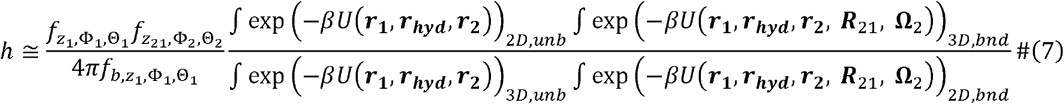

where the length scale coefficients *f* capture the 2D restraints and are defined by:

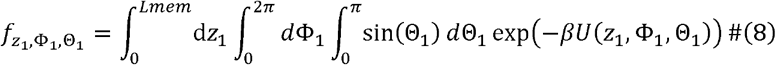

Where the subscript *b* refers to the bound dimer. The integral *dz*_1_ is over *Lmem* to indicate that not all values of *z*_1_ are allowed for the proteins to remain adhered to the membrane. We do not need to specify *a priori* the value of *Lmem*, but the fluctuations of the protein in *z*_1_ when bound to the membrane will set the range. If the protein dissociates from the membrane, *z*_1_> *Lmem*, those configurations should not contribute to the configuration integrals for bound or unbound ensembles. To further simplify Eq. 7, we assert that the configuration integral of the unbound state is the same in 2D and 3D, and we assert the bound state is the same in 2D and 3D. If we further assume that the potential energy is separable in the 2D height, pitch and roll, *U*(z_1_,Φ_1_,Θ_1_), =*U* (z_1_) +*U*(Φ_1_) +*U* (Θ), then Eq. 8 becomes

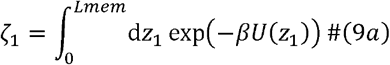

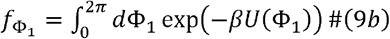

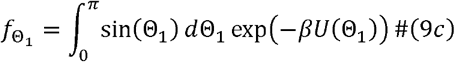

And Eq. 7 becomes

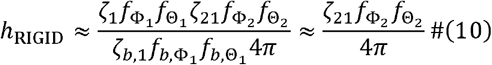

Eq. 10 is the same as the result found in previous work^18, 43^ that characterized 2D binding for rigid domains, where the focus was on binding between adhesion receptors across a cell-cell membrane gap, rather than the peripheral membrane proteins studied here. The values in Eq.9 will be less than the 3D values due to the membrane, 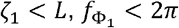 and 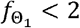. If the membrane gap, rather than the peripheral membrane proteins studied here. The values in Eq. restraints on protein 1 are comparable in the unbound and bound ensemble, which is feasible for two proteins on the same membrane, then we get the final expression on the right in Eq. 10. In this rigid-body limit, the 2D relative to 3D dissociation constant is the lengthscale that is determined only by the *z* fluctuations when membrane bound, multiplied by the fractional loss in pitch and roll angles.

### II. C Violations of the rigid-body approximation

Proteins are not truly rigid, and in II.B above we made three major assumptions to reach Eq. 10 from Eq. 7: A) there is no change in configurational entropy at the backbone or residue level from 3D to 2D. B) there is no change to the (relative) potential energy of sampled configurations from 3D to 2D. C) the configurations that are lost from 3D to 2D due to restrictions to height, roll, and pitch have potential energies that make a negligible impact on the bound state partition to more strongly favor (*h<h*_*rigid*_) *or* disfavor (*h<h*_*rigid*_) 2D association. function. Violations to these assumptions can cause deviations from the rigid-body predictions to more strongly favor (*h* < *h*_rigid_) or disfavor (*h* > *h*_rigid_) 2D association.

Violations that would further stabilize 2D association (*h<h*_*rigid*_ as we see for LSP_M3.0) could either be driven by a loss of configurational entropy in the unbound ensemble (A), and/or a lower potential energy of the bound ensemble for configurations sampled in 2D (B). Regarding the first point, for proteins that form a significant interface with the membrane, the inherent flexibility of the backbone and residues could be reduced or altered via surface localization, beyond the rigid-body restrictions we have accounted for already. Given a matching rigid-body orientation, a loss in flexibility would always reduce the range over which the integrals are evaluated in the 2D state, such that the scalar ratio in Eq. 7 is decreased:

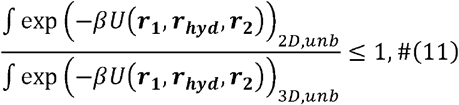

assuming that the potential energy *U* across equivalent degrees of freedom is unchanged. However, we note that if the same loss of flexibility occurred in the bound ensemble, it would have the opposite effect on the scalar ratio, acting to de-stabilize 2D association, as

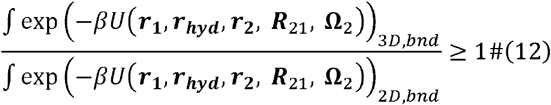

again assuming the same potential energies. What we see below for LSP_M3.0 is instead primarily due to (B): the internal configurations sampled in 2D are not identical to those sampled in 3D, due to the intrinsic flexibility of the proteins, and they have a lower potential energy in the bound ensemble, thus stabilizing 2D association by a larger denominator in Eq. 12. More generally, if the protein undergoes a conformational change upon membrane binding in either bound or unbound states, it could sample different potential energies, and neither of the inequalities in Eq. 11 or 12 need hold.

Violations that could de-stabilize 2D association (*h > h*_*rigid*)_ could be driven by A, B, or C. If loss of flexibility in 2D is higher in the bound state with no other changes, Eq. 12 will increase the scalar ratio. For (C), we know for a fact that the configurational space is distinct for the integrals in Eq. 12, as both *R*_21_and Ω_2_ have reduced sampling in 2D via Z_21_,Θ_2_,Φ _2_. We increase the scalar ratio. assumed they were comparable if the bound state contains a narrow set of 2D accessible structures with sufficiently low potential energy. Then the Boltzmann weight of those configurations will dominate, rendering all other configurations, including those ‘lost’ in 2D, negligible, such that ⎰ exp(-*βU* (r_1_r_*hyd*_,r_2_,R_21_, Ω_2_)) _3*D,bnd*_ ≈ exp(-*βU*_*min)*._ In the extreme example, 2D restrictions could eliminate the most stable bound structures, thus dramatically increasing above the rigid body prediction.

## III. RESULTS

### III. A For the LSP_M3.0 dimer, the crystal state structures are much more stable in the 2D environments

We constructed the free energy surface (FES) for a pair of LSP1 monomers in three distinct environments: solution (3D), bound to a lipid membrane bilayer (2D), and restricted on a 2D ‘pseudo-membrane’ (2D-Ps) while fully solvated in water (Fig 1). We perform the same calculations for LSP_M3.0 (simulated with MARTINI 3 forcefield, Movie S1-S3) and for LSP_M2.2p (MARTINI 2 forcefield version 2.2P, Movie S3-S6), noting that the results from LSP_M2.2p in 3D solution were published previously^16^. The FES was constructed using enhanced sampling via metadynamics (Fig 2, Methods). In all three environments, our sampling produced many transitions between the bound and unbound states, and at least 2 or more transitions to the crystal structure, indicating good sampling of configuration space (Fig. 2g,h and Fig S2). We have compared the surface across multiple time blocks to assess the confidence of our FES features and shown that the bound ensemble relative to the unbound ensemble are robust enough to give significant estimates for Δ*G* (Fig S3, Fig S4, Fig S5). Our confidence of our FES features and shown that the bound ensemble relative to the unbound previous work showed that the LSP_M2.2p monomers in solution (3D) engage in several nonspecific bound dimer states in addition to the known crystal structure^16^. We similarly find here that LSP_M3.0 in solution samples diverse nonspecific states. However, only for the LSP_M3.0 dimer do we see that localization to the membrane effectively eliminates these nonspecific dimers, leaving only the stable native dimer.

**Figure 2.**
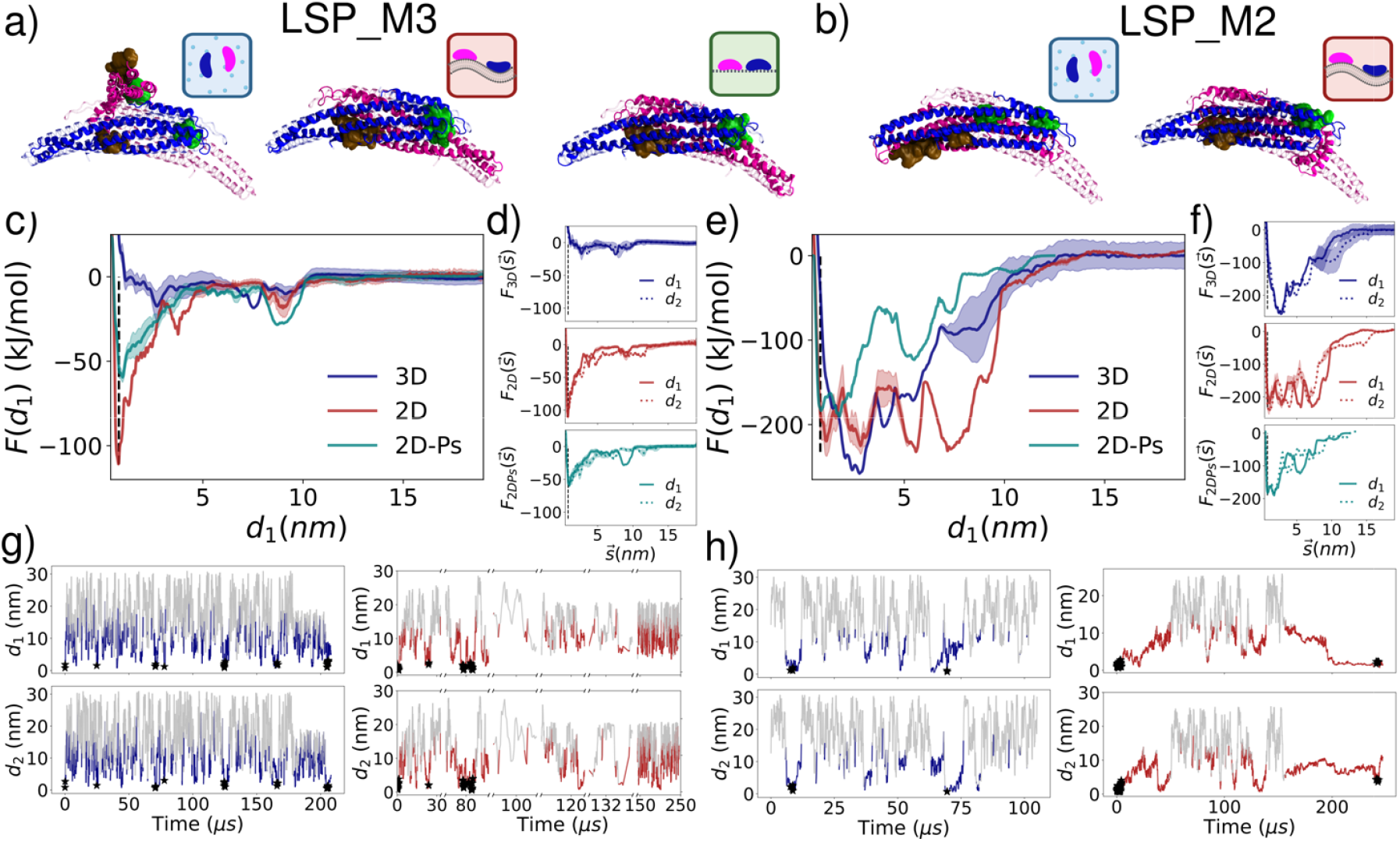
The FES reveals significant changes to the 2D bound ensemble compared to 3D for LSP_M3.0. The left half of the figure shows the FES results from LSP_M3.0, and the right half from LSP_M2.2p. **a-b)** From the FES for each environment, we show the most stable dimer structure corresponding to the minimum free energy. The atomistic representation of the two monomers in dark blue and magenta is obtained from backmapping the coarse-grained structures for **a)** LSP_M3.0 and **b)** LSP_M2.2p. The dimer is represented with the blue monomer aligned to the crystal structure. The light pink outline shows the position of the pink monomer in the crystal structure. The LSP_M3.0 in 2D selects for dimers very similar to the crystal state, unlike 3D. The LSP_M2.2p dimers are slightly shifted relative to the crystal state. **c)** 1-dimensional Free Energy surface (FES) vs d_1_, obtained by integrating the free energy over d_2_, shows that the crystal structure (denoted by dashed black line) is the most stable state in 2D membrane and 2D-Ps. We show the error estimates in light colors around the sample mean by considering a 95% confidence interval (Methods). The most stable state in 3D corresponds to a non-specifically bound dimer state as seen in a). **d)** 1-dimensional FES for each environment show a similar dependence on either d_1_ (solid) or d_2_ (dashed). **e)** 1-dimensional FES for LSP_M2.2p simulations shows broad and extraordinarily deep energy wells for both 3D and 2D simulations. The most stable state in 2D is closer to the crystal structure than in 3D. However, there are several stable non-specifically bound states in both environments. **f)** 1-dimensional FES for each environment is similar vs d_1_ (solid) or d_2_ (dashed). **g)** To demonstrate sampling, the time evolution of the CVs d_1_ (top panel) and d_2_ (bottom panel) is plotted every 100 ns 3D (blue) and 2D (red) for LSP_M3.0. There are multiple transitions from the unbound state (gray; d_1_, d_2_ > 13.0 nm) to the bound state (colored; d_1_ and d_2_ < 13.0 nm) with some of them also sampling the crystal state (denoted by black stars; dRMSD < 2.0 nm). **h)** Similar plots for LSP_M2.2p simulations are shown for 3D (blue) and 2D (red). The LSP_M2.2p simulations observe multiple transitions from the bound to unbound state but fewer samples of the crystal state. We note that the monomers in the 2D LSP_M3.0 simulations sometimes dissociated from the membrane (or adhered to the opposite membrane leaflet). These time points represented by disjointed segments in the x-axis are excluded from the analysis as they do not sample the 2D bound or unbound ensemble (33–66 µs, 95.2-97.8 µs, 101.9-103.3 µs, 127.6-128.5 µs and 138-141 µs).

For the LSP_M3.0 dimer, the FES shows that the bound states, and particularly the crystal structure configuration, are significantly more favorable in the 2D membrane and pseudo-membrane environments than in 3D (Fig. 1 and 2a,c,d). Integration of the corresponding FES indicates that free energy of dimerization is much lower in 2D than in 3D (Table 1, Fig S5). A significant distinction between the 3D and 2D environments is that the native crystal state in the 3D environment is not the most stable substructure of the bound ensemble and does not form the native dimer interface. In contrast, for the 2D and 2D-Ps FES, the most stable structure is very close to the crystal structure. The FES generated from our 2D pseudo membrane simulations is in fact remarkably similar to the full bilayer 2D simulations, much more so than to the 3D system (Fig 1, Fig 2c), despite retaining only the rigid-body geometric restraints of the 2D environment via the monomer’s position and two orientational angles (Methods, Fig S6-S11, Movie S3). These restraints are sufficient to stabilize the crystal state of the LSP1 dimer relative to any other bound states, unlike in 3D. The 2D-Ps FES is not quite as deeply in favor of the crystal structure compared to the 2D system with the explicit bilayer (Fig 2a), which we revisit below.

**Table 1.** Free Energy differences (in kJ/mol) for dimerization in different systems (see Methods).

|  | LSP_M3.0 |  |  | LSP_M2.2p |  |
| --- | --- | --- | --- | --- | --- |
|  | 3D | 2D | 2D-Ps | 3D | 2D |
| (kJ/mol) | $-21.2 \pm 7.9$ | $-98.7 \pm 6.9$ | $-62.3 \pm 5.2$ | $-250.9 \pm 29$ | $-228.9 \pm 12$ |
|  | 38,800 | 1130 | 1130 | 54,900 | 1130 |

### III. B For the LSP_M2.2p dimer, in contrast, the dimer does not benefit significantly from 2D localization

Unlike the LSP_M3.0 dimer, the FES of the LSP_M2.2p dimer shows comparable overall stability between the bound and unbound states in both 3D and 2D (Fig 2). For LSP_M2.2p, the 3D and 2D FES both favor a variety of nonspecific bound structures, producing a broad and deep energy well in the bound states, albeit more funnel-shaped towards the lowest energy structure in 3D. Somewhat surprisingly then for LSP_M2.2p, the restriction to the membrane surface does not increase the selectivity for a single bound state, in contrast to LSP_M3.0. The most stable structures for LSP_M2.2p in both 3D and 2D show deviations from the crystal structure (Fig 2b). The second monomer is translated along the native dimer interface in 3D, creating an almost completely overlapping interface between the two monomers. This compresses the dimer length and disrupts the characteristic crescent shape of the BAR domain dimer. The native dimer interface is better preserved in the most stable structure of the 2D environment, but the monomer tends to fold back on itself at one end, again increasing the protein-protein interface and disrupting one side of the crescent-shaped dimer. The pseudo-membrane simulations produce a similarly deep energy well with a stable structure that is between the 2D and 3D structures (Fig 2e,f), but the statistics are not sufficient to confidently assess the uniqueness of its shape.

While the major qualitative difference between LSP_M2.2p compared to LSP_M3.0 is that the 2D environment does not improve LSP_M2.2p’s selectivity or stability for a specific, near-native bound state, there are also clear quantitative differences between the two dimers. The parameterization of the LSP_M2.2p dimer produces extraordinarily favorable potential energies of bound states and ultimately binding free energies Δ*G* that are not representative of The parameterization of the LSP_M2.2p dimer produces extraordinarily favorable potential standard protein-protein interactions as seen in ours^16^ and other work^45^. These parameters promote large desolvation barriers for dimers^40^ and enhanced stickiness that drives frequent non-native dimer interfaces^46, 47^, as we see here. The newer LSP_M3.0 parameterization has much improved accuracy in binding energies^38, 39^ and we therefore expect it to be a more accurate model of the ‘true’ experimental system^22, 24^. However, our study is designed to quantify relative behavior and is thus equally applicable to both dimers.

### III. Rigid-body approximation *h*_RIGID_ predicts sub-nanometer lengthscale from observed fluctuations

We here estimate *h*_RIGID_ if our proteins were purely rigid bodies, as defined in Eq. 10. We unbound therefore assume these rigid proteins retain the same (narrow) set of specifically bound structures with the same potential energy difference Δ*U* between the bound and ensembles in both 3D and 2D. The predicted *h*_RIGID_ is thus purely entropic and determined by fluctuations of the membrane bound monomers around their z-position, *z*_disp,_ pitch Θ and roll Φ angles. The observed distributions from the MD simulations of these three variables (Fig 3) show significant restrictions in the 2D and 2D-Ps systems compared to 3D, whereas the yaw orientation is unconstrained as expected in all systems (Fig S12). By numerically integrating (Methods) over the product of distributions in Fig 3 (Eq. 9), for LSP_M3.0, 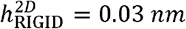, and 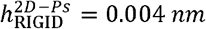. Both these lengthscales are smaller than in previous studies which reached down to ∽ 0.1*nm*^18^. As anticipated, this is due to the proximity of the BAR domain’s dimer interface to the membrane surface and the extended protein-membrane interface that reduce translational and orientational degrees of freedom. The pseudo membrane value is smaller than 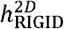 because its fluctuations in *z* are more narrowly distributed (Fig 3c), stemming from our constraint potential being derived based on a minimal *z* separation between the protein and the membrane (Methods, Fig S13). The *z*_*disp*_ distance more accurately captures fluctuations due to both membrane and protein, and this larger variation in *z* -sampling for the 2D vs 2D-Ps simulations (Fig S13) could help explain the lower free energies observed on the bilayer (Fig 2).

**Figure 3.**
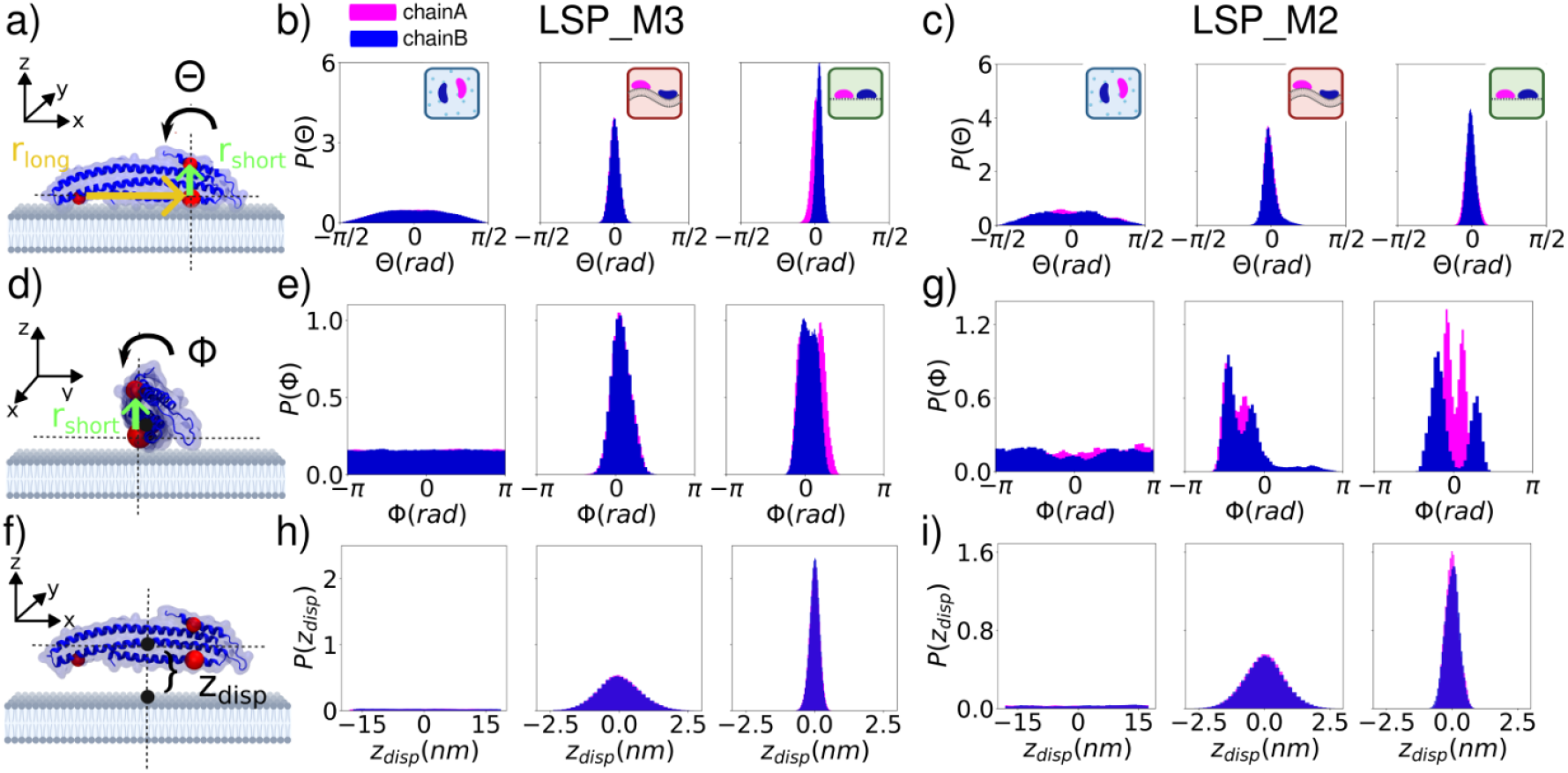
Sampled rigid-body degrees of freedom in three environments for LSP_M3.0 and LSP_M2.2p quantify effects of 2D restriction. Top row: **a)** illustration of the pitch angle for a monomer. Pitch angle distributions for **b) LSP_M3.0 c) LSP_M2.2p**. Histogram of the pitch angle () shows a uniform distribution in 3D space such that. All distributions are plotted with the sample mean subtracted off. The distribution on the membrane and pseudo-membrane shows restricted sampling as expected, with values of chain A in pink and chain B in blue throughout. **Middle row d)** illustration of the roll angle for a monomer **e**,**f)** The roll angle () also shows uniform sampling when in solution such that, but restricted sampling in the 2D simulations. **Bottom row g)** illustration of the fluctuations in, the distance of the monomer centers-of mass from the origin (3D), or membrane (2D). **h**,**i)** is sampled uniformly in 3D as expected. The 2D values are restricted close to the membrane, and the pseudo membrane is more tightly restricted because of our restraint definition. In all cases, data was collected throughout the metadynamics simulations, histogrammed, and normalized.

For LSP_M2.2p, we find similar values and trends, with 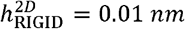 and 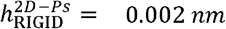, again due to the more restrictive z-constraint potential applied for the pseudo membrane. Unlike the other dimer, for LSP_M2.2p, we find that the range of pitch angles sampled can vary significantly throughout the entire bound ensemble (Fig 3d), deviating from the narrow set of values sampled for LSP_M3.0 (Fig 3b), or observed in the native dimer state used to construct the pseudo membrane constraint (Fig S9-S10). Our analysis shows that the stable non-specific dimers on the membrane are driving this broader range of pitch angles for LSP_M2.2p that deviate from angles sampled by the native dimer (Fig S14).

### III. Rigid-body approximations cannot explain observed change in binding free energies from 3D to 2D

To compare the rigid-body estimates *h*_RIGID_ with the values constructed directly from the sampled FES using Eq. 5, we evaluated the Δ*G* values for each environment and system using block averaging (Methods, Fig S5, Table 1). Because fluctuations and uncertainty in will exponentially impact our calculation of from Eq. 5 (Table S1), we here directly compare the distribution of values from the FES (collected over the last half of the sampling) to the predicted values, which we compute by inserting Eq. 10 into Eq. 5 to get:

This rigid body prediction for is also a distribution due to its dependence on the collected from the 3D FES simulations. With our calculated values in the 0.002-0.03 nm range, this offset term amounts to kJ/mol, so mean values can also be estimated from Table 1.

In Figure 4, we show that the predicted values deviate significantly from the sampled values for LSP_M3.0 (2D and 2D-Ps) and for LSP_M2.2p (2D). We performed Kolmogorov-Smirnov (KS) tests to confirm that the values are not sampled from the same distribution as the FES values, and therefore the rigid-body approximation fails. We verified this conclusion holds even with 50% fewer data points included from the simulations (). For LSP_M3.0, the rigid-body estimate is too high, and the simulations sample lower free energies on the membrane but also the pseudo-membrane, or. For LSP_M2.2p, in contrast, the rigid-body estimate is lower than the sampled values, indicating that the membrane acts to destabilize the bound state compared to a purely entropic, rigid-body prediction,. Both configurational entropy and enthalpy could drive either of these trends (see section II.C).

**Figure 4.**
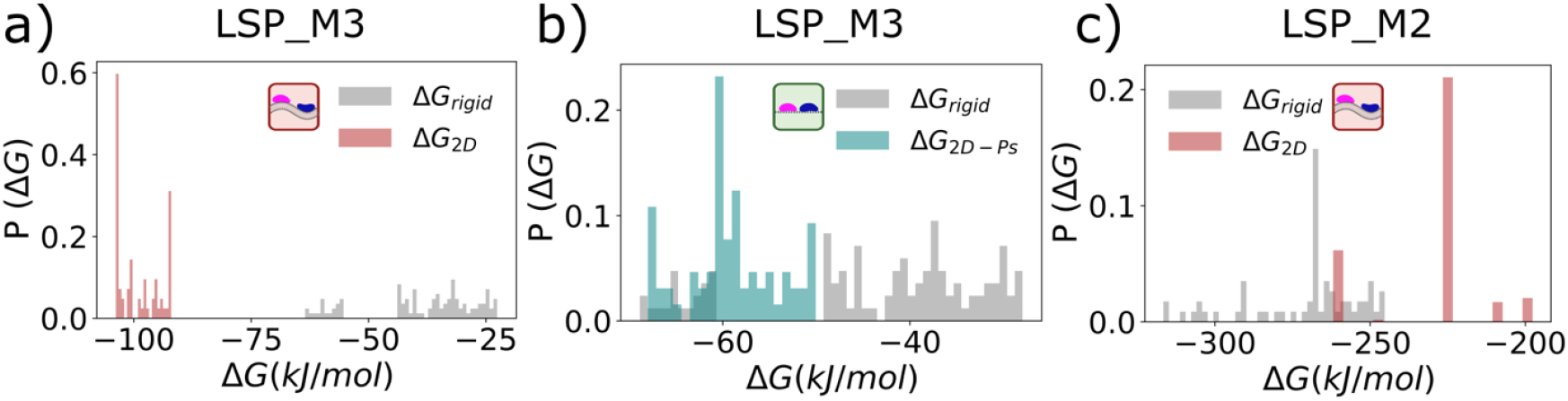
Hypothesis testing of the rigid-body approximation shows that it fails to predict the observed 2D free energies. a) The binding free energies calculated from the FES (red bars) are compared to the rigid-body predictions (gray bars) from the 3D FES values and Eq. 13 for LSP_M3.0. The FES values are stored every 1 s from the last 50% of the simulation to provide a distribution in both 2D and 3D of 70-100 points for all panels. LSP_M3.0 samples lower free energies on the membrane that are not consistent with the rigid body predictions (KS test 10^−38^), and b) the same for the pseudo membrane, with the FES values from simulation in green, and in gray (KS test 10^−25^). c) For LSP_M2.2p, the sampled free energies in 2D are actually higher than the rigid body prediction and are not consistent with them (KS test 10^−34^). These results are robustly preserved with 50% fewer data points in the distributions. For LSP_M3.0, the bilayer free energies (2D) are also statistically distinct from the pseudo membrane (2D-Ps) (KS test 10^−33^).

### III. D. Configurational sampling is distinct in 2D compared to 3D but does not fully explain differences in *h* estimates

To go beyond a rigid-body comparison of the sampling in 3D versus 2D and consider all internal degrees of freedom, we assessed the fluctuations of each monomer relative to fixed rigid-body orientations. For LSP_M3.0, we find that the monomers in 2D adopt configurations that are significantly closer to the crystal structure than the 3D monomers (Fig 5a). Because we have aligned all monomers to the same reference (crystal) structure, these variations can only emerge due to the flexibility of the backbone and side chain beads. The measured fluctuations across the backbone beads corroborate this, showing for example dampened fluctuations of the membrane-binding residues compared to 3D (Fig 5b). Notably, these configurational changes are not induced by dimerization, as they are retained even for the unbound monomers, indicating it is the surface restriction that selects for these configurations. Remarkably, the 2D and 2D-Ps results are nearly perfectly overlapping to one another, indicating that neither explicit contacts with lipids nor the charged environment of the membrane surface are necessary to shift these configurations, but only the rigid-body restrictions imposed by the 2D surface. For the LSP_M2.2p, we see quite different behavior; the 2D and 2D-Ps configurations are both centered around the same RMSD=0.4nm value as the 3D configurations in both the bound and unbound states, showing that membrane restriction is not helping select for more native-like configurations (Fig 5c). For the bound state (Fig 5c), the distributions are not identical, with the 3D monomers sampling a long tail of larger RMSD values and the 2D monomers populating a

**Figure 5.**
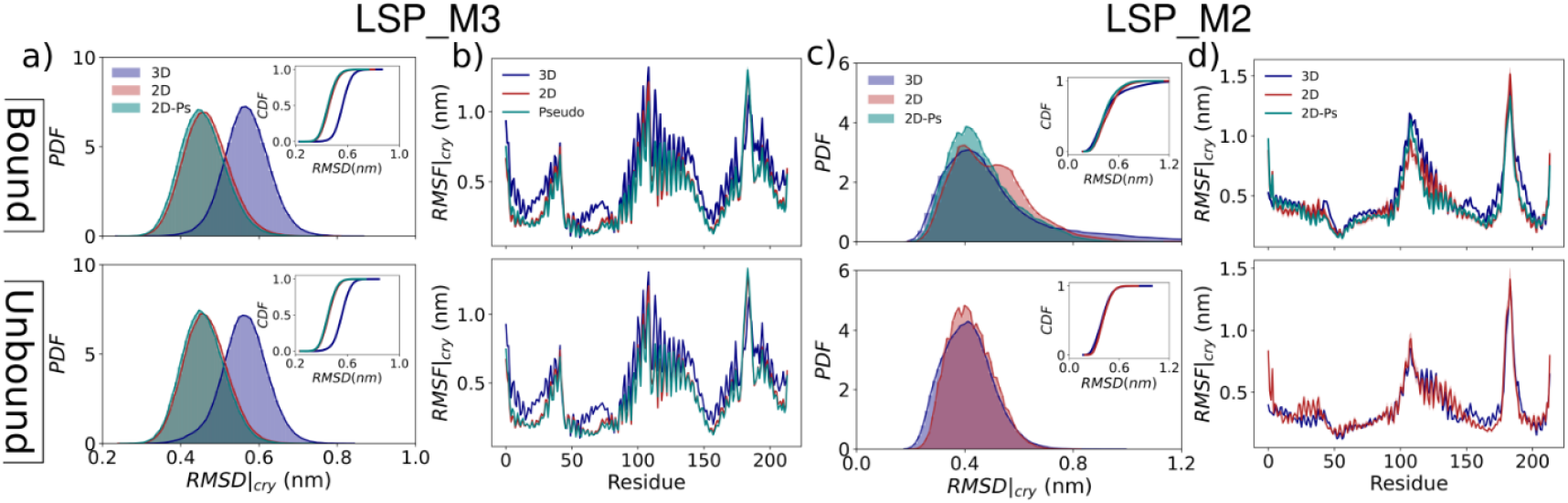
Configurational sampling and flexibility of LSP1 monomer varies between different environments. **a-b) LSP_M3.0. a)** The RMSD for each protein monomer is evaluated relative to the crystal structure monomer following alignment (SI)^48^. We separate monomers in the bound ensemble (top) from the unbound ensemble (bottom) although they are nearly indistinguishable for LSP_M3.0. The 2D (red) and 2D-Ps (green) distributions are overlapping one another but clearly distinct from the 3D (blue) distribution. The CDF (inset) also emphasizes the stark contrast in conformations sampled between 2D and 3D. **b**) The root mean square fluctuations (RMSF) of each residue in the monomer are also evaluated relative to the crystal structure, reinforcing the smaller deviations in the 2D and 2D-Ps environments compared to 3D, including stretches containing membrane-binding residues 56ARG, 63LYS, 66LYS, 70ARG, 126ARG, 130LYS, 133ARG. **c-d) LSP_M2.2p c)** RMSD and **d)** RMSF measurements show substantial overlap between 2D, 2D-Ps and 3D, with all three distributions centered around the same RMSD value, unlike for LSP_M3.0. The unbound monomers (bottom) are in relatively close agreement in both 3D and 2D. The RMSF measurements show that the LSP_M2.2p protein has a similar trend in flexibility across the length of the protein compared to LSP_M3.0, with peaks in the same locations, but with better conservation across all three environments.

shoulder at RMSD=0.5 nm. In both cases these larger variations are stemming from the strong nonspecific contacts in the bound ensemble that can induce bending and twisting of the monomers (Fig 2b), which we quantified as producing a markedly larger distribution of end-to-end distances for LSP_M2.2p (Fig S15). Unlike for LSP_M3.0, we see here that these flexible states are induced by binding and are thus largely absent from the unbound ensemble (Fig 5c). The RMSF across the LSP_M2.2p backbone atoms are broadly similar to the trends in LSP_M3.0, with peak local flexibilities around residues 110 and 180. Consistent with the RMSD, they are more similar across all 3 environments.

These observed differences between environments demonstrate configurational sampling that is not captured by the rigid-body approximation to *h*_*RIGID*_. For LSP_M3.0, we showed above that *h* ≪*h*_*RIGID*_ and in section II.C we justify how this can occur if the number of these internal configurational states in the 2D unbound ensemble is reduced relative to 3D, without any change to potential energy. Although the RMSD distributions in Fig 5a are only 1D projections and cannot thus report on the full configurational entropy, we note that their highly similar variances suggest that the degree of internal sampling or number of states has not changed dramatically from 3D to 2D. Next we will thus assess whether the clear changes to the types of configurations has therefore produced a change in enthalpy, via sampled potential energies.

### III. The relative potential energies are not conserved for bound structures from 3D to 2D environments

We show here that the shift in sampled configurations from 3D to the 2D environments (Fig 5) is accompanied by changes to the relative potential energies Δ*U*_LJ_ of the bound structures (Fig 6). We show here that the shift in sampled configurations from 3D to the 2D environments (Fig 5) is The clear trends observed for LSP_M3.0 structures shifting more towards the native configuration in 2D and 2D-Ps (Fig 5a) correlate with an increased stability of the non-bonded protein-protein Lennard-Jones (LJ) interaction in 2D and 2D-Ps as the dimers more closely approach the native crystal state (Fig 6a). A similar trend is observed for the LJ protein-water interaction but in reverse—as the proteins approach the dimer state in 2D, the protein-water interaction is destabilized relative to the 3D structure (Fig 6a). Because these contributions to Δ*U*_LJ_ are not the same in 3D versus 2D, this also violates the assumptions of our rigid-body approximations made from Eq. 7 to Eq. 10. The other potential energy terms including the Coulombic interactions show similar trends, but with a smaller magnitude (Fig S16). For LSP_M2.2p, we again see distinct behavior from LSP_M3.0, as we anticipated based on the overlapping configurational sampling in all three environments (Fig 5). The Δ*U*_*LJ*_ values from 3D to 2D to 2D-Ps criss-cross one another as the dimer RMSD (dRMSD, see SI) approaches the native dimer structure for both protein-protein and protein-water interactions (Fig 6b). While there are significant differences that emerge, which are likely due to the large variety of stable non-specific structures in the bound ensemble (Fig 2), the 2D structures are not consistently more stable than the 3D structures. For structures at Drmsd ≤ 2 nm, nm, the 3D structures typically have a more stable protein-protein Δ*U*_LJ_ and this could be contributing to our finding that for LSP_M2.2p, *h* ≪*h*_*RIGID*_.

**Figure 6.**
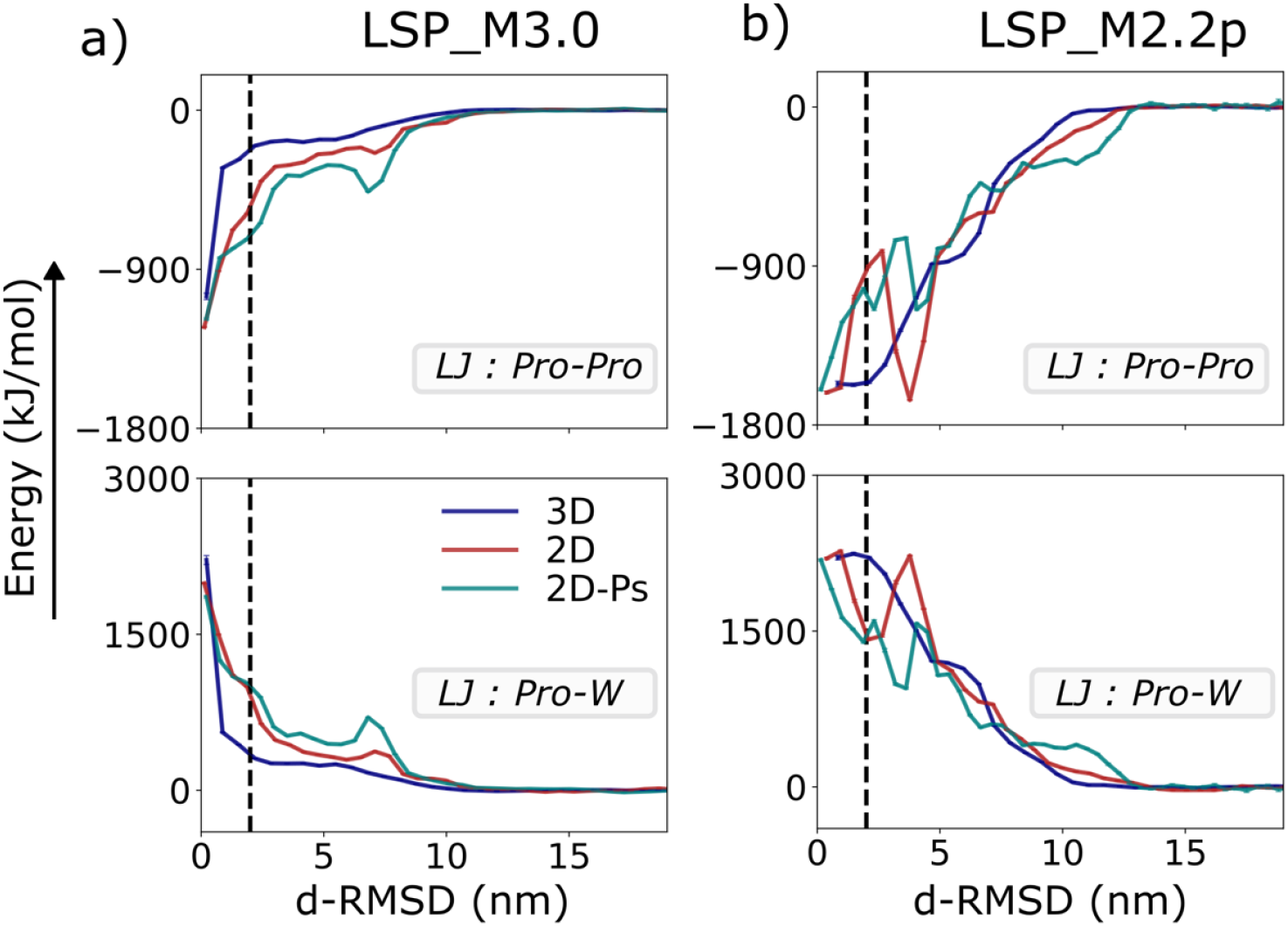
Potential energies from non-bonded contacts varies across environments for bound states. a) LSP_M3.0. b) LSP_M2.2p. The potential energy due to the Lennard-Jones (LJ) potential interaction between the two proteins (top Pro-Pro) and between the protein beads and water (bottom: Pro-W) are evaluated for configurational ensembles that are sorted by dimer RMSD (dRMSD). The black dashed line indicates dRMSD = 2 nm, a useful cutoff for structures highly similar to the native crystal dimer. Large dRMSD capture unbound states. We zero-out the LJ potential in all panels relative to the value in the unbound state. As expected, all environments show a plateau in the LJ energy for the unbound states (dRMSD>12nm). **a)** For LSP_M3.0, the 2D environments have significantly stronger protein-protein LJ energies that stabilize configurations in the bound ensemble. The 2D environments have weaker protein-water LJ energies in 2D compared to 3D, in reverse of the protein-protein trend. **b)** For LSP_M2.2p, the LJ energies do not show a clear trend with dRMSD, likely due to the abundance of nonspecifically bound states, although they do show significant differences across the three environments. Each dRMSD bin has error bars (calculated as SEM across number of configurations ∼ 500) which are less than the linewidth.

The rigid-body assumption demands that the two proteins maintain the same binding interface during dimerization in the two different environments. The potential energies indicate this is not the case (Fig 6). We performed a deeper inspection of the absolute potential energies of the near-native dimers, including structures with i) dRMSD < 0.6 nm (1.0 nm for LSP_M2.2p) and ii) 0.6< dRMSD < 2.0 nm. For LSP_M3.0, the 2D environment selected for these near-native structures more frequently in both classes and with lower average potential energies for the 2.0nm class (Fig 7a, Fig S17). For the 3D structures, the larger gain in protein-water energy moving from the 0.6nm class to the 2.0nm class makes the 2.0nm class more relatively favorable than the 0.6nm class. In 2D, this is not the case. Visualizing the ensemble of aligned dimer structures via trajectory snapshots reveals that while the 3D and 2D structures are quite similar for the 0.6nm class, there is much more variety in the 2.0nm 3D structures (Fig 7c) and the distribution of dRMSD values is skewed towards larger values in 3D compared to 2D (Fig S17). Several of these structures are accessible in 3D but not with the 2D restrictions to rigid-body sampling, thus the 2D environment has eliminated near-native structures that have less favorable potential energies. The pseudo membrane results are more aligned with the 2D energetics, consistent with their overlapping monomer configurations (Fig 5), albeit with enough variations in potential energy to provide some support for the overall less stable binding free energy (Fig S18). For LSP_M2.2p, as we now expect, the 2D environment does not show a consistent improvement in selectivity of near-native dimers compared to 3D (Fig 7b). The 1.0nm class is more frequently observed in 2D, but the 2.0nm class is not. The energetics of the 2.0nm class overlaps and even can exceed the stability of the 1.0nm class, reinforcing the poor selectivity of the LSP_M2.2p system for the native dimer structure over competing dimers (Fig 7b). The trajectory snapshots of LSP_M2.2p (Fig 7d) show a larger conformational diversity for 3D configurations at 2.0nm, which is similar to LSP_M3.0. However, the 2D structures for both 1.0 and 2.0nm classes of LSP_M2.2p show the end distortion of the dimer interface, illustrating again that the explicit bilayer in LSP_M2.2p is not as effective at selecting for the native dimer interface.

**Figure 7.**
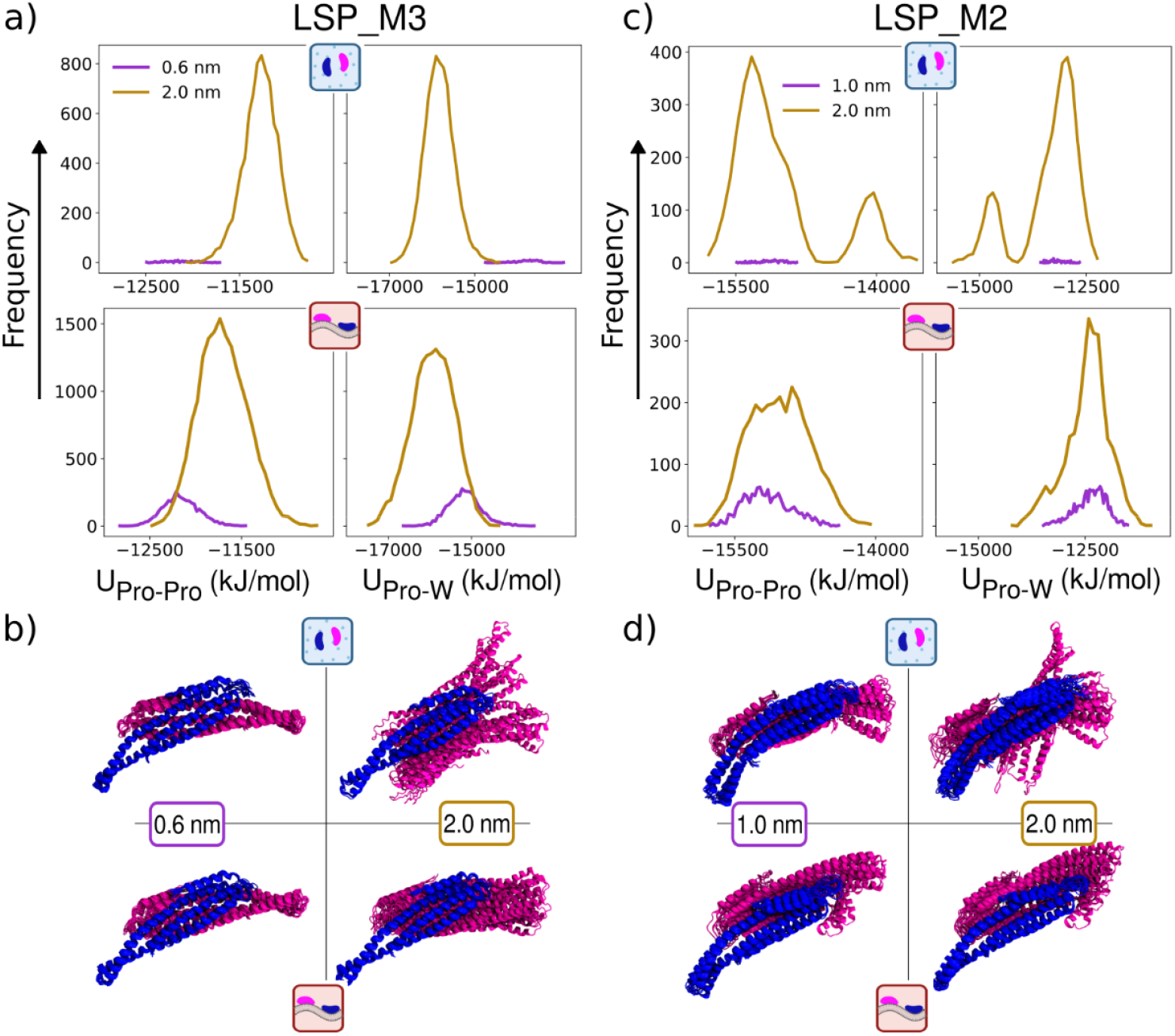
The near-native dimer structures are sampled more commonly in 2D and can produce lower potential energies for LSP_M3.0. The distribution of LJ potential energies for near-native bound dimer structures that have a dRMSD < 0.6 nm (purple curves) or 0.6< dRMSD < 2.0 nm (gold curves). **a) LSP_M3.0** From 3D (top row) to 2D (bottom row), the protein-protein energetics (left column) span a similar range of values for 0.6nm (purple), but are more frequent in 2D, given ∼350,000 frames of bound configurations from each environment. The 2.0nm structures are more stable in 2D than in 3D and more frequent. The protein-water energetics in the right column. **b)** Combined trajectory snapshots of the LSP_M3.0 dimer in the two configurational states (left vs right) and in 3D (top) vs 2D (bottom). **c) LSP_M2.2p**. 2.0nm structures in 3D sample two states with different energies. The more stable state has similar energetics to the 0.6nm states and is closer to the native crystal dimer. Although the 1.0nm states are more frequently sampled in 2D, the 2.0nm states are more frequently sampled in 3D. **d)** Combined trajectory snapshots for LSP_M2.2p show slight variations in the 1.0nm structures at the flexible end. The 2.0nm structures show higher variability in 3D compared to 2D, as in (b).

## IV. DISCUSSION

A key finding of our work is that dimerization on the membrane can diverge from a rigid-body approximation to significantly improve (LSP_M3.0: *h* ≪*h*_*RIGID*_) or diminish (LSP_M2.2p: A *h* ≪*h*_*RIGID*_ the selectivity and stability of the bound state. The BAR-domain dimer we studied here represents an important test case of the rigid-body approximation, as these proteins reside in the 3D cytoplasm but localize to membranes for their primary function. Because the BAR-domain dimer interface is very close to the membrane surface, unlike previously studied adhesion protein dimers^17, 18, 29^, one might anticipate that the enthalpy or internal configurational entropy of the binding process could change in 2D, as we found here. Critically, for the more experimentally accurate parameterization^38, 39^ for the LSP_M3.0 dimer, our results show that the restrictions of the proteins to the 2D surface dramatically improves their selection of the bound state, which aligns with their evolutionary purpose of assembling on membranes and we discuss further below. Our computed lengthscale for is well below the nanometer lengthscale, meaning that for BAR domains that reside in a cell with a 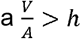,which is all cell types, binding will be favored in 2D with a maximal gain to the affinity given by 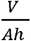 ^26^. Given our computed 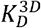 from the average Δ *G* in the 0.01 *μM* regime (Table 1 and Eq. 1), the 3D system will already have mostly LSP1 dimers (concentration is ∼1*μM* from UniProt), but 2D localization will then push all monomers into the dimer state. We predict that the membrane localization is more critically primed for triggering BAR domain oligomerization, which is expected to have a significantly weaker 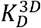 than the dimerization and is observed to produce limited complexes in the cytoplasm^24^. Surprisingly, the improved selectivity of the 2D proteins for the bound state did not require an explicit lipid bilayer, as this same trend was reproduced in our pseudo membrane simulations that recapitulated only the rotational and translational restrictions of the surface on the protein motion. The explicit membrane does improve the selectivity and stability even further beyond the pseudo membrane, which is likely due to the enhanced motion in *z* accessible on the fluctuating membrane compared to our tighter constraints on the pseudo membrane (Fig 3). These gains on the membrane occurred despite that the BAR domain did not undergo large conformational changes, as can happen for unstructured proteins^37^.

How transferrable or generalizable are our results? The rigid body approximation *h*_*RIGID*_ provides a useful baseline from which to consider how enthalpy or configurational changes in 2D will further modify the value. MD simulations (or Monte Carlo) therefore offer a critical tool to first calculate *h*_*RIGID*_ from the fluctuations used for Eq. 10. These same MD simulations can then further help estimate its accuracy prior to the more time-consuming free energy calculations performed here needed to evaluate the true (Eq. 5). Our results for the two dimers LSP_M3.0 and LSP_M2.2p were distinct due to the variations in the force-field, but we could rationalize the free energy trends from 3D to 2D at least partially by a relatively straightforward analysis of the configurations sampled in each environment relative to a rigid-body reference state (the crystal structure), and by comparing the potential energies across each environment. For LSP_M3.0, both 2D environments drive a marked shift in the conformations of the LSP1 monomers relative to 3D (Fig 5), and a corresponding lowering of potential energies of near-native bound structures (Fig 6-7). For LSP_M2.2p, in contrast, while the 2D environments produce some deviations in the conformations sampled relative to 3D, there is no systematic ‘improvement’ in conformations or the corresponding potential energies. While these are imperfect metrics for free energy, they both demonstrate that the rigid-body approximation is violated and enthalpy does play a role. For LSP_M3.0 these metrics correctly indicated that *h*_*RIGID*_ is an over-estimate of *h*. As we stated above, it evolutionarily makes sense that the membrane environment at least ensures a value of *h* in the nanometer regime that promotes more dimerization on the surface. However, this implies that the LSP_M2.2p results make little evolutionary sense, as *h* is in the micron regime (Table S1). We attribute this to the unphysically stable interactions of this force-field that favor protein association broadly with reduced selectivity^16, 40, 45^. In absolute terms, the LSP_M2.2p dimer state will be close to irreversible due to the free energy barrier for unbinding in both 3D and in 2D 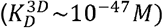 hence neither environment would observe monomers at any physiologic concentration. In other words, membrane localization cannot trigger dimerization that is already guaranteed in 3D. A natural follow-up to this study would be to use all-atom simulations to quantify the BAR domain FES with higher accuracy compared to experiment^49^. While these simulations are far more expensive, a faster alternative to metadynamics that is particularly logical for a 3D to 2D comparison would be free energy calculations using a cascade of geometric restraints, as they directly operate on Euler angles and translational degrees of freedom, with good agreement to experiment^44, 49, 50^.

While we do not have experimental estimates of *h* for BAR domain proteins, we argue that our results are consistent with in vitro experiments and that other BAR domain dimers likely produce comparable behavior. Measurements of BAR-domain dimerization including LSP1 are convoluted by the fact that the dimers can then form higher-order oligomers^14 22, 51^, which introduces a distinct (and weaker^10, 36^) protein-protein interface. Nonetheless, experiments find significant increases in oligomerization (and thus dimerization) on membranes compared to in solution for a variety of BAR proteins ^14, 52-54^. At least one of these binding interfaces therefore favors 2D assembly via dimensional reduction, or 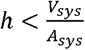.For many BAR domains we also expect minimal dimers to form in the cytoplasm on their own, based on experimentally measured 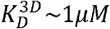 ^9, 10, 36^ and the more than 10x lower concentrations than this in HeLa cells (endophilin 1 and FCHO2) ^55^. These same BAR-domain proteins clearly form assemblies on membranes^14, 24^, and while additional protein interactions could certainly play a role for these often multi-domain proteins^56^, the assembly appears biased towards 2D. The LSP1 BAR domain has even more extreme assembly behavior in 2D, forming large oligomeric scaffolds called eisosomes containing thousands of LSP1 copies on the plasma membrane^23^, with only small cytoplasmic complexes observed^24^.

Overall, this simulation and theoretical approach provides mechanistic insight into how a key peripheral membrane protein class, the BAR domain, has evolved to ensure dimerization is more favorable on the membrane surface. Extension to other peripheral protein interactions would help establish how broadly 2D localization is exploited to trigger self-assembly. For the BAR domain, this approach could also be extended to assess the strength of the oligomerization contacts between BAR domains on the membrane, or any curvature-dependence of its interactions, as these proteins are known to cause tubulation and sculpting of membranes in vitro and in silico^53, 57, 58^, even driving membrane fission^59^. While these domains seem to bind or induce curvature in a manner dependent on the membrane curvature ^60, 61^, their 2D affinities might also benefit from a more curved membrane. Here we studied dimerization only on a flat membrane, but any mechanical or curvature dependencies that could impact 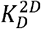 could further lower (or increase) *h* thus shifting the equilibrium population of dimers and oligomers^26^. Finally, these lengthscales *h* represent essential parameters in kinetic and reaction-diffusion models that necessarily integrate out the detailed structural information present here to be able to predict population-level dynamics as they occur in living systems^62, 63^. In this capacity, not only the equilibrium lengthscales but also the kinetic comparison 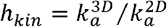,where 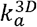 are the association rate constants, are important. Extending rigid-body studies of rates^64^ to again consider conformational or enthalpic effects in peripheral proteins could similar enhance our understanding of 2D timescales, which experimentally seem to support *h*_*kin*_ values also in the nanometer scale^31, 33, 65^. Evaluation of rate ‘constants’ in 2D requires more care, as unlike in 3D, they are not truly constant due to reentrant diffusion, which complicates their estimation^66^. Understanding the physico-chemical determinants of these fundamental 2D vs 3D binding metrics is critical to accurately predict how any surface localization that supports 2D diffusion of binding partners will control when and where self-assembly occurs.

## V. METHODS

### V. A Molecular Dynamics Simulations

*System Setup:* We performed MD simulations with three different system environments, (a) solvated and freely diffusing proteins, (b) proteins bound to a membrane and (c) fully solvated proteins with translational and rotational motion restricted to a 2D ‘pseudo’ membrane (see Fig 1). All three environments were simulated using the newly parametrized MARTINI 3.0 forcefield ^39^ and using the MARTINI 2.0 forcefield, using the GROMACS package^67^ (version 2021.4 for MARTINI 3.0 and 2020.2 for MARTINI 2.0). All systems were simulated using metadynamics (aside from equilibration and a few short unbiased trajectories) using the PLUMED plugin (version 2.8.0)^21^. The structure for LSP1 was obtained from the PDB database (accession code: 3PLT) ^22^. Two copies of chain A were used for the homodimer since chain B had missing residues. The LSP1 homodimer was coarse grained with the ELNEDIN elastic network (spring force constant 500 kJ mol^-1^ nm^-2^ and cutoff as 0.9nm) using the *martinize*.*py* tool downloaded with the open beta version 3.0.b.3.2, followed by correction of side chain dihedrals as recommended using the *bbsc*.*sh* script.

For the solution simulations, the initial configuration of the two monomers are in their bound state, as extracted from the crystal structure. The proteins are solvated using the standard water model with 0.1M of NaCl in a simulation box of ∼ 36 × 36 × 36 nm^3^ under periodic boundary conditions. Energy minimization of systems was followed by 5 ns of equilibration in the NVT ensemble and an additional 5 ns of equilibration in the NPT ensemble for our MARTINI 3.0 models and as previously published for our solution MARTINI 2.0 simulations^16^. The relatively large box size was used to ensure that for these dimerization simulation studies, proteins oriented in an end-to-end fashion would not be able to simultaneously interact with their physical partner at one end and the periodic image of the same protein at their other end, as that would mimic a filament and not a dimer.

For membrane simulations, the LSP1 dimer was added to an equilibrated bilayer in a box of size ∼ 38 × 38 × 19 nm^3^.The phospholipid bilayer was setup using the *insane*.*py* tool (modified for MARTINI 3.0 lipids) with a composition of 80% POPC, 10% POPS and 10% POP6 (MARTINI for PIP_2_). The bilayer was first neutralized and solvated with the standard water model and 0.1 M NaCl followed by a similar protocol of minimization and equilibration runs. The LSP1 monomers in the bound dimer form (like seen in the PDB structure) were then added to the equilibrated bilayer and positioned such that its membrane binding residues face the membrane. After adding the protein, the system was again minimized and equilibrated following the same protocol. For membrane simulations with MARTINI 2.0 we followed these same steps from the beginning, except swapping out the force-field topology and molecule parameter file (bead names).

For the pseudo-membrane systems we use a simulation volume of ∼ 37.0 × 37.0 × 37.0 nm^3^. The system was set up using a configuration of the two MARTINI LSP1 monomers present in the bound state, at the end of a short unbiased simulation on the membrane surface. The dimer was then isolated and solvated with the standard water model and 0.1 M NaCl. First the system was energy minimized, then equilibrated using 5ns NVT, followed by 5ns NPT. We then introduced our harmonic restraint potentials to both monomers in a stepwise manner. The three harmonic restraint potentials over the height z_surf_ and orientations α_long_, α_short_ each monomer are detailed below. They were switched on in the order *U*(z_surf_), *U*(α_long_), *U*(α _short_ over 1400ns and kept on for the remainder of the simulations. To switch them on, we started off with a low or ‘soft’ force constant, letting it adjust to each restriction before ramping up the force constant to its final target value. The simulation parameters were otherwise the same as the other system environments.

#### Simulation parameters

All simulations were performed using improved MARTINI parameters^68^.

The electrostatic interactions were modeled using the Reaction-field method with a 1.1 nm cutoff. The dielectric constant was set to default value of 15 with an 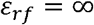,The Lennard-Jones potential was used with a ‘vdw-modifier’ to smoothly shift to zero after a cutoff of 1.1 nm. We used the leapfrog integrator with a timestep of 20 fs. Neighbor lists were constructed using the Verlet scheme with nstlist set to 10 and the Verlet-buffer-tolerance (VBT) was kept at the default value of 0.005 kJ mol ^−1^ ps^-1^. However, these default parameters were shown to contribute to artifacts in the pressure tensor leading to unphysical undulations in the membrane as reported by^69^. While we did not observe any unphysical characteristics in our system, we altered the VBT parameter to 0.0002 kJ mol ^−1^ ps^-1^ after 156 µs for MARTINI 3.0, as recommended. The temperature was maintained at 310 K using the velocity rescaling thermostat while the pressure was maintained at 1 bar (semi-isotropic for membrane simulations) using a Berendsen barostat for equilibration (coupling constant of 5.0 bar^-1^) and a Parinello-Rahman barostat of 12.0 bar^−1^ for the production runs.

#### Establishing constraints for a 2D pseudo membrane system

In our pseudo membrane simulations, the LSP1 monomers are simulated fully water-solvated in a 3D environment, without any membrane. However, of the 3 translational and 3 rotational rigid-body orientational axes for rotation, we introduce constraints on 3 of them: translational motion in the z-direction, and rotational motion around the long and short axis of each monomer. Translational motion in x and y are unconstrained, as is rotation about the z-axis, or the effective yaw angle, since both an proteins and lipids diffuse. We parameterized harmonic constraints (SI Methods) on each of the three variables z_surf,_ α_long_, α_short_ by quantifying the observed values that are sampled during an unbiased simulation of almost 1 µs when our proteins are bound to an actual membrane (Fig S6, S7, S8 for LSP_M3.0, and Figures S9, S10, S11 for LSP_M2.2p). The monomers were bound into the dimer state for the full trajectory, although statistics collected on unbound monomers were not substantially different. We defined z_surf_ = z_*patch*_ ± z_*Mpatch*_ .The position on the protein, ***r***_*patch*_ = [*x*_*patch*_, *y*_*patch*_, z_*patch*_] is defined via the center of geometry of all residues that participate in the membrane binding patch, 56ARG,63LYS,66LYS,70ARG,126ARG, 130LYS,133ARG. The corresponding membrane patch ***r***_***Mpatch***_ = [*x*_*Mpatch*_, *y*_*Mpatch*_, z_*Mpatch*_]is defined by all lipid headgroups within 1.2 nm from any of these protein residues.

To quantify the sampling of the orientations of each monomer, we define a vector 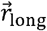 connecting beads 84LYS to 150 THR along the long-axis of the monomer. We measure the orientation of this vector relative to the global z-axis,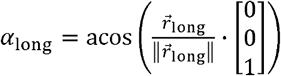.To quantify thesampling relative to another nearly orthogonal axis, we define a vector 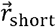 connecting beads 259GLU to 80ARG along the short-axis of the monomer, pointing ‘up’ if the protein is bound to the membrane. We again compute the vector orientation relative to the global z-axis via 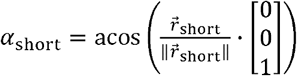 These angles are straightforward to calculate for performing restraints during our pseudo membrane simulations. However, we note that for theoretical calculations of Eq. 14 and in the analysis section below, we define the calculation of Euler angles for our monomers. The distributions for all three variables are fit to harmonic potentials (SI Methods) with parameters reported in SI (Table S2, Table S3).

#### Metadynamics Parameters

For building the free energy surface (FES), we primarily used the standard metadynamics technique ^70^, and in some cases we switched on (and off) the well-tempered bias ^71^ to help with convergence (SI Methods). The FES was evaluated with respect to two collective variables (CVs), *d*_1_ and *d*_2_ representing distances between two pairs of sites that are in contact in the bound crystal structure, as established in our previous work^16^. Both variables are illustrated in Fig 1. The first collective variable *d*_1_is the distance between the center of geometry of two sites comprised of residues 57LYS-64THR on chain A and residues 78GLU-92ASP on chain B. The second collective variable *d*_2_ is the distance between the sites consisting of residues 224LEU-229ASP on chain A and residues 189ASN-196LYS.

We collect all metadynamics parameters in Table S4 for all three system environments. Following our previous study^16^, the deposition times τ_*G*_ for each bias potential were determined based on the relaxation time of our CVs to travel the distance σ_1_,σ_2_ that define the widths of the the widths σ_1_,σ_2_, to retain a similar deposition time (SI Methods). bias potentials. Because sampling is slower for proteins on the membrane system, we reduced the widths σ_1_,σ_2_, to retain a similar deposition time (SI Methods).

*Wall potential:* To accelerate sampling in our two CVs *d*_1_ and *d*_2_, we introduced a restraint potential using PLUMED:

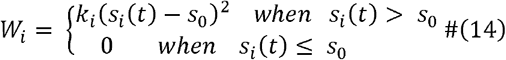

where *i* =1,2 corresponding to *d*_1_ and *d*_2_, k_*i*_ =500 kJ mol^-1^ nm^-2^, and *s*0 =21.0 nm for the solution simulations and 19.0 nm for the 2D simulations. We could not simply reduce the box-size because this could introduce artifacts from a monomer interacting with a partner and its periodic image.

### V. B Numerical analysis of MD simulations

*Numerical evaluation of binding free energy* Δ *from simulations:* From our metadynamics simulations, we compute the free energy surface *F*(*d*_1_, *d*_2_). This FES is related to the canonical partition function *Z* (*d*_1_, *d*_2_) via 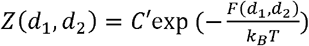, where is a constant ^20^. To compute the relative free energy of the bound to the unbound state, we must integrate over this

partition function for values of *d*_1_, *d*_2_ that reside in the bound ensemble vs those in the unbound ensemble. We use a cutoff *rcut* in both values of the CV to partition the surface into the bound with a large enough value of *rcut* Smaller cutoff values can leave low-energy states in the and unbound ensemble. We demonstrate in Fig S19 that the free energy difference converges unbound ensemble; due to the Boltzmann weighting, this misrepresents the relative unbound state free energy. We use numerical integration over both CVs via

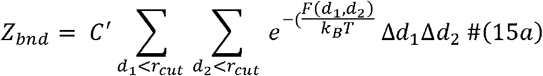

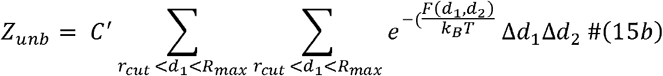

where *R*_*max*_ is the maximal separation of the CV in our system, which is limited by the restraining wall we use to either *R*_*max*_ =21*nm* (3D) or 19 *nm* (2D), or 36 *nm* for LSP_M2.2p in over when computing configurations that contribute to the partition function, we use 3D. This establishes the maximal volume of the unbound ensemble V is the volume sampled over when computing configurations that contribute to the partition function, we use 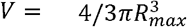, and in 2D 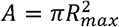, The partition functions in Eq. 15 are input into Eq. 6, then Eq 1 or Eq 7 to compute Δ*G* and *K*D.

*Evaluating convergence and error of the FES and binding free energy*: To compare the free energy differences Δ*G* between different systems and obtain error estimates, we performed block averaging. The free energy difference was calculated as the mean Δ*G* over *N* blocks where the first block consists of Δ*G* evaluated at 50% completion of the total simulation time and the subsequent 50% of the simulation divided into *N*-1 blocks. We show progression of FES at 50%, 80%, 90%, 95% of the total time to illustrate that despite fluctuations in some regions of estimates ε on our sample mean Δ*G* by considering a 95% CI by using the following phase space, the relative depths is reasonably preserved (Fig S3, S4). We evaluated the error estimates ε on our sample mean ⟨Δ*G*⟩ by considering a 95% CI by using the following expression^72^ 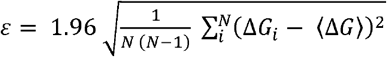 where the 1.96 comes from computing the z-score of a normal distribution at 95%. We evaluated the error by considering different block sizes (varying *N*) for each simulation as shown in Fig S5, showing good convergence. For the final free energy estimates, we used a block size of 20 µs for 3D MARTINI 3 and MARTINI 2 simulations. For all 2D simulations a larger block size of 40 µs was used. These estimates are reported in Table 1.

#### Measuring Euler/Tait-Bryan angles for our monomers

Given the long and short axes of our monomer *m* at any point in time, 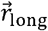 and 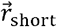, we can define a right-hand orthogonal frame with unit vectors where 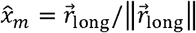 is the so-called roll axis, 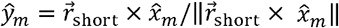 is the pitch axis, and 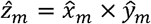 is the yaw axis. We follow the Tait-Bryan angles for frame z, y’, and x’’, where primes indicate rotated frames. We can thus compute yaw 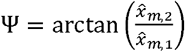, pitch 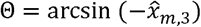 and roll 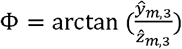, intrinsic rotations around the global where the subscript indicates the vector index. We computed distributions of sampled values for these angles in both the bound and unbound ensembles. This is necessary to compute *h*RIGD from Eq. 10.

#### Measuring the height variations of membrane bound proteins

To quantify the height fluctuations of our monomers for Eq. 14, we measured Z _disp_ = Z _*cogp*_ – Z _*lipid*_. The protein height is determined by the monomer center-of-geometry *r*_*cogp*_ = [*x*_*cogp*_, *y* _*cogp*_, *z*_*cogp*_]. The height displacement uses z _*lipid*_, the coordinate of a single lipid located at the same position as *x*_*cogp*_, *y* _*cogp*_ at that time point. We calculate z *disp* for the lipid on the outer and the inner leaflet, for comparison, with similar results (Fig S13).

#### Visualization of structures

The cartoon representations for the proteins were obtained by backmapping the corresponding coarse-grained structures using the backmap tool ^73^. Images and movies were renderes using ^74, 75^.

*Numerical evaluation of rigid-body approximation, h* RIGID *from simulation data:* From Eq 10, we can quantify the ratio of 2D to 3D dissociation constants by assuming our proteins are rigid, and the roll variable, we computed *U*(Φ_1_) numerically by histogramming all observed Φ_1_ values, measuring the change in sampling of the roll, pitch, and height variables on the membrane. For the roll variable, we computed *U*(Φ_1_) numerically by histogramming all observed Φ_1_ values, normalizing to a probability distribution, and using *U* (Φ_1_) =-*k*B *T* ln (*p*(Φ)), with enforced (*U* (Φ_1_))= 0. The same was done for the pitch and height (Fig 3). These distributions are then input into Eq. 9 and numerically integrated to calculate, 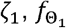, and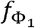, for each unbound monomers, a standard deviation of *h*RIGID for the membrane systems is ∼0.001nm, and for the monomer and for each monomer in the bound dimer. Based on the variance between the bound monomers, a standard deviation of **h**_**RIGID**_ for the membrane systems is ∼0.001nm, and for the 2D-Ps systems is ∼0.0003nm, or 10% in both cases.

## SUPPORTING INFORMATION

- Supplementary Information (pdf): SI Methods including extended theoretical derivation, SI Figures S1-S19, and SI Tables S1-S4.
- Supplementary Movies 1-6 (mp4): Trajectories of LSP1 simulations in all 3 environments for LSP_M3.0 and LSP_M2.2p.

## AUTHOR INFORMATION NOTES

## Corresponding Author

Margaret E. Johnson

## ACKNOWLEDGEMENTS

M.E.J. gratefully acknowledges funding from a National Institutes of Health MIRA Award R35GM133644. This work was carried out at the Advanced Research Computing at Hopkins (ARCH) core facility (rockfish.jhu.edu), which is supported by the National Science Foundation (NSF) grant number OAC 1920103. This work also used Bridges-2^76^ at Pittsburgh Supercomputing Center through allocation MCB150059 from the Advanced Cyberinfrastructure Coordination Ecosystem: Services & Support (ACCESS) program, which is supported by National Science Foundation grants #2138259, #2138286, #2138307, #2137603, and #2138296. We thank Dr Alexander Sodt for helping us setup the membrane simulations and Johnson Lab members for feedback.

---

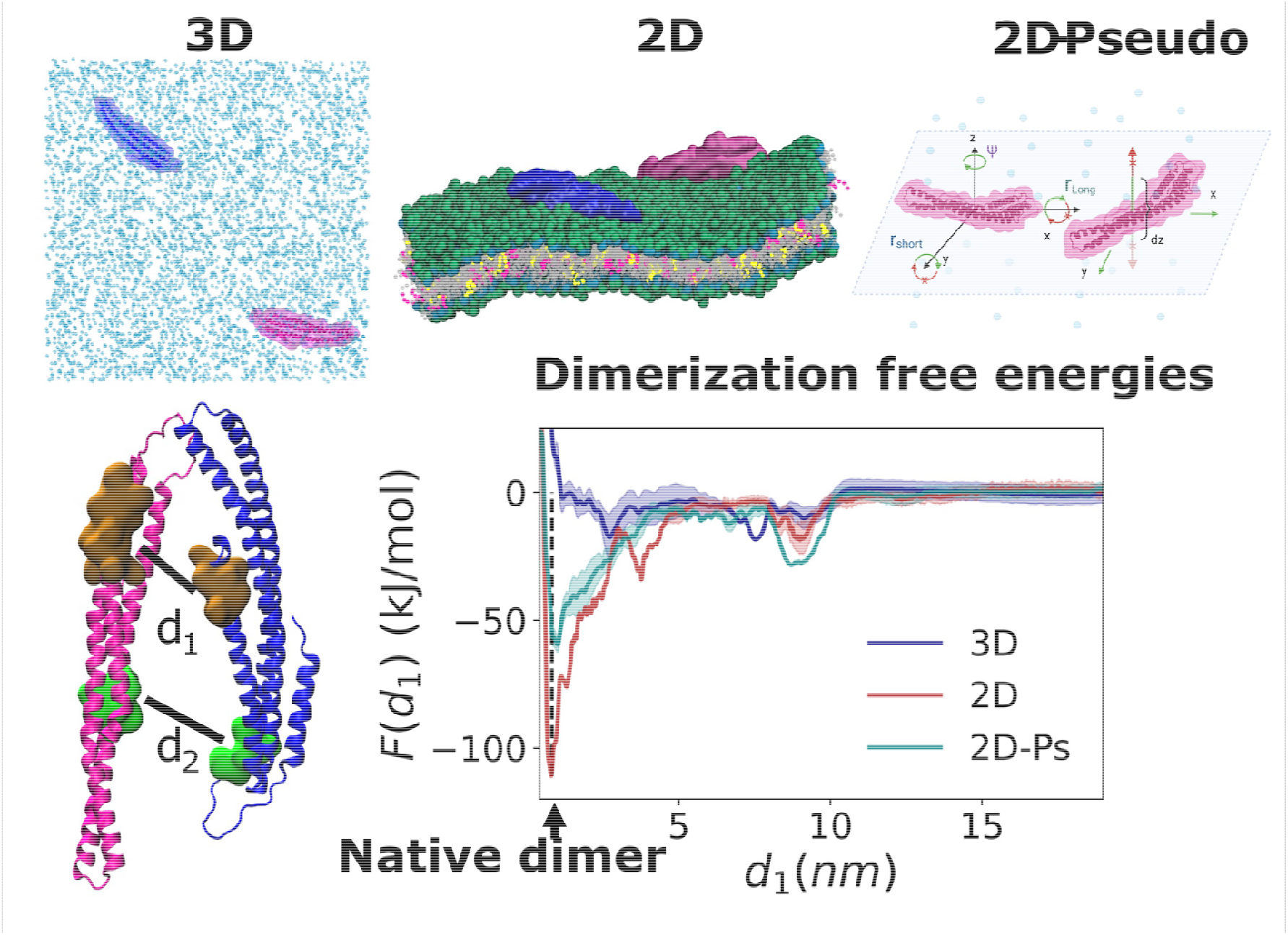
For Graphical Table of Contents Only: BAR domain dimerization free energies are calculated in solution (3D), on a membrane bilayer (2D), and in a pseudo membrane restraint simulation (2D-Ps). The 2D environments both select for much more stable native dimer free energies (green and red curves) compared to 3D (blue curve). Analysis reveals backbone changes are responsible for improving the dimer enthalpy in 2D, stabilizing the bound state.

