## Supplemental Information for "Binding affinities for 2D protein dimerization benefit from enthalpic stabilization"

**for**

### SUPPLEMENTAL METHODS

#### SI. Extended theory background from Main section II.A

For binding in 3D, it is well-established that by definition the binding free energy scales with volume as $\Delta G(V_1) = \Delta G(V_2) + k_{\mathrm{B}}T\ln\left(\frac{V_{2,bnd}V_{1,unb}^2}{V_{1,bnd}V_{2,unb}^2}\right)$, and simplifies when the volume is the same in the bound and unbound ensemble, $V_{bnd} \cong V_{unb}$, to:

$$\Delta G(V_2) \cong \Delta G(V_1) + k_{\mathrm{B}}T\ln\left(\frac{V_2}{V_1}\right) \qquad (S1)$$

The binding free energy is destabilized with increased volume in 3D. This is because larger volume stabilizes the unbound state more than the bound state. We report $\Delta G$ values for our sampled system $V$, which defines the same $K_D$ as the standard state via Eq. 1 of the main text.

In 2D, the relative free energy for surface-restricted binding instead scales as

$$\Delta G(SA_2) \cong \Delta G(SA_1) + k_{\mathrm{B}}T\ln\left(\frac{SA_2}{SA_1}\right) \qquad (S2)$$

where we assumed $SA_{bnd} = SA_{unb}$ for both volumes. Increases to the height of the system volume should not affect the binding free energy of 2D associated partners, as the additional volume is not sampled by either the bound or unbound monomers.

The Helmholtz free energy relates to the canonical partition function via $F = -k_BT\ln(Q)$. The (dimensionless) partition function is defined for a system of $N$ particles using,

$$Q = \frac{1}{n_S! h^{3N}} Z_p(\{\boldsymbol{p}^N\}) Z(\{\boldsymbol{r}^N\}) \qquad (S3)$$

where $h$ is here Planck's constant and there will be an indistinguishability factor for (at least) the $n_S$ solvent particles. $Z_p(\{\boldsymbol{p}^N\})$ is the momentum integral, which we assert will remain the same value for bound and unbound ensembles, given the fixed temperature. $Z(\{\boldsymbol{r}^N\})$ is the configuration integral dependent on positions $\boldsymbol{r}$ of all $N$ atoms, which differs 1) between bound and unbound ensembles, and 2) as membrane localization restricts protein positions in the volume. The ratio of the bound and unbound partition functions $Q_{bnd}/Q_{unb}$ will thus reduce to the ratio of the bound and unbound configuration integrals:

$$\Delta F(V) = -\, k_B T \ln\left(\frac{Z_{bnd}(\{\boldsymbol{r}^N\})}{Z_{unb}(\{\boldsymbol{r}^N\})}\right) \qquad (S4)$$

Which is the same as Eq. 3 of the main text.

To expand the partition functions explicitly, we use similar notation to Ref[1] and [2] where the coordinate system illustrated in Fig S1 defines the $N_1$ internal coordinates of protein 1 $(\boldsymbol{r_1})$, protein 2 $(\boldsymbol{r_2})$ , of the $N_s$ solvent particles $(\boldsymbol{r_w})$, and of membrane coordinates $(\boldsymbol{r_m})$ expressed relative to the translational positions of proteins 1 and 2 in the system volume. We will simplify this notation further to $\boldsymbol{r_1}, \boldsymbol{r_2}, \boldsymbol{r_{w,m}}$ for compactness. The translational position and rigid-body orientation of protein 1 in the volume are given by $\boldsymbol{R}_1$, $\boldsymbol{\Omega}_1$ respectively. The relative position of protein 2 with respect to 1 is then given by $\boldsymbol{R}_{21}$ and its rigid-body orientation given by $\boldsymbol{\Omega}_2$. To describe the total partition function, we separate configurational space into bound and unbound states so the total configuration integral is given by $Z = Z_{bnd} + Z_{unb}$. To classify configurations into bound or unbound, we use the Heaviside function $H_s$, with the unbound ensemble selected via $H(|\,\boldsymbol{R}_{21}| - r_{cut})$ and the bound ensemble via $H(r_{cut} - |\,\boldsymbol{R}_{21}|)$. We use $r_{cut}$ as a distance threshold to distinguish the bound dimer from two non-interacting monomers. This threshold may generally be dependent on the internal coordinates of both monomers, their hydration solvent, and their relative orientations, $r_{cut} = r_{cut}(\,x_{21}, y_{21}, z_{21}, \boldsymbol{\Omega}_2, \boldsymbol{r_1}, \boldsymbol{r_2}, \boldsymbol{r_{w,m}})$, with this volume then defined as $V_{cut}$.

The configuration integrals can then be written to separate out the integrals over the position/orientation of protein 1 $(x_1, y_1, z_1)$ that each span a boxlength $L$, and $\boldsymbol{\Omega}_1$ the rigid-body orientation of molecule 1 at a given position using the Euler/Tait-Bryan convention with $\Psi_1$ the yaw, $\Theta_1$ the pitch, and $\Phi_1$ the roll as defined (Methods) and illustrated in Fig S1. The integrals prior to any integration are written:

$Z_{unb} = \int_V \, d\,\boldsymbol{R}_1 \int d\,\boldsymbol{\Omega}_1 \int_V \, d\,\boldsymbol{R}_{21} \int d\,\boldsymbol{\Omega}_2 \int d\boldsymbol{r_1} d\boldsymbol{r_2}\, d\boldsymbol{r_{w,m}} H(|\,\boldsymbol{R}_{21}| - r_{cut}) \exp\left(-\beta U\left(\boldsymbol{R}_1, \boldsymbol{\Omega}_1, \boldsymbol{r_1}, \boldsymbol{r_2}, \boldsymbol{r_{w,m}}, \boldsymbol{R}_{21}, \boldsymbol{\Omega}_2\right)\right)$ (S5a)

$Z_{bnd} = \int_V \, d\,\boldsymbol{R}_1 \int d\,\boldsymbol{\Omega}_1 \int_V d\,\boldsymbol{R}_{21} \int d\,\boldsymbol{\Omega}_2 \int d\boldsymbol{r_1} d\boldsymbol{r_2}\, d\boldsymbol{r_{w,m}} H(r_{cut} - |\,\boldsymbol{R}_{21}|) \exp\left(-\beta U\left(\boldsymbol{R}_1, \boldsymbol{\Omega}_1, \boldsymbol{r_1}, \boldsymbol{r_2}, \boldsymbol{r_{w,m}}, \boldsymbol{R}_{21}, \boldsymbol{\Omega}_2\right)\right)$ (S5b)

Where $\beta = 1/k_B T$. We rewrite these integrals to denote what integrals can be performed immediately in 3D (but not fully in 2D), where $s \in (bnd, unb)$:

$$Z_s = \int_0^L \mathrm{d}z_1 \int_0^L dy_1 \int_0^L dx_1 \int_0^{2\pi} d\Psi_1 \int_0^{2\pi} d\Phi_1 \int_0^{\pi} \sin(\Theta_1)\, d\Theta_1 Z_{21,s} \qquad (S6a)$$

$$Z_{21,s} = \int_V d\,\boldsymbol{R}_{21} \int d\,\boldsymbol{\Omega}_2 \int d\boldsymbol{r_1} d\boldsymbol{r_2}\, d\boldsymbol{r_{w,m}} H_s \exp\left(-\beta U\left(\boldsymbol{R}_1, \boldsymbol{\Omega}_1, \boldsymbol{r_1}, \boldsymbol{r_2}, \boldsymbol{r_{w,m}}, \boldsymbol{R}_{21}, \boldsymbol{\Omega}_2\right)\right) \qquad (S6b)$$

Where $Z_{21,s}$ contains all the remaining integrals that define the relative configurations between the two monomers and their internal coordinates.

For a protein 1 that is free to diffuse in all 3 dimensions, the system's potential energy does not depend on its translational position or rigid body orientation within the volume (e.g. there are no boundary effects). Thus, we can simplify the potential energy function $U$ (Eq. S6b) to $U\left(\boldsymbol{r_1}, \boldsymbol{r_2}, \boldsymbol{r_{w,m}}, \boldsymbol{R}_{21}, \boldsymbol{\Omega}_2\right)$ and integrate:

$$Z_s^{3D} = 8\pi^2 V\, Z_{21,s}^{3D} \qquad (S7)$$

Following this integration over protein 1's position and orientation, we can now consider its central position and orientation fixed in the box, and the integrals over $\boldsymbol{R}_{21}$ and $\boldsymbol{\Omega}_2$ thus occur over $x_{21}, y_{21}, z_{21}$, and $\Psi_2$ the yaw, $\Theta_2$ the pitch, and $\Phi_2$ the roll of protein 2.

Returning to Eq. S6, the 2D system differs in several key respects. Both proteins 1 and 2 are bound to the membrane surface, and we specify the membrane oriented in the x-y plane such that its normal typically points in the z-direction (Fig 1 main text). The integrals over the translational position and orientation of protein 1 (and 2) in the volume are now restricted by the membrane in height $z_1$, the pitch $\Theta_1$, and the roll $\Phi_1$. The other three integrals over $x_1, y_1, \Psi_1$ are unrestrained, and similar to 3D we reduce the the potential energy function $U$ to $U\left(z_1, \Phi_1, \Theta_1, \boldsymbol{r_1}, \boldsymbol{r_2}, \boldsymbol{r_{s,m}}, \boldsymbol{R}_{21}, \boldsymbol{\Omega}_2\right)$. These integrals can then be immediately integrated in Eq. S6a to produce:

$$Z_s^{2D} = 2\pi A \int_0^{Lmem} \mathrm{d}z_1 \int_0^{2\pi} d\Phi_1 \int_0^{\pi} \sin(\Theta_1)\, d\Theta_1 Z_{21,s}^{2D} \qquad (S8)$$

where $A = L^2$. The integral $\mathrm{d}z_1$ over $Lmem$ indicates that not all values of $z_1$ are allowed for the proteins to remain adhered to the membrane. In 2D, $r_{cut}$ defines the $A_{cut}$ for the bound ensemble, as the height $Lmem$ is restricted for both bound and unbound ensembles.

### SII. Derivation of the simplified $h$ ratio of Main text Eq. 7

To simplify the configuration integrals for Eq. S6b, we proceed with simplifications that each assume the potential energy can be decoupled for subsets of variables.

1. The potential energy of the unbound ensemble is independent of the relative separation $\boldsymbol{R}_{21}$ and orientation $\boldsymbol{\Omega}_2$ of protein two relative to one, as the monomers are no longer interacting: in 3D: $U(\boldsymbol{r_1}, \boldsymbol{r_2}, \boldsymbol{r_{s,m}}, \boldsymbol{R}_{21}, \boldsymbol{\Omega}_2) = U(\boldsymbol{r_1}, \boldsymbol{r_2}, \boldsymbol{r_{s,m}})$. In 2D we retain the dependence on height, pitch, and roll: $U(z_1, \Phi_1, \Theta_1, \boldsymbol{r_1}, \boldsymbol{r_2}, \boldsymbol{r_{s,m}}, \boldsymbol{R}_{21}, \boldsymbol{\Omega}_2) = U(z_1, \Phi_1, \Theta_1, \boldsymbol{r_1}, \boldsymbol{r_2}, \boldsymbol{r_{s,m}}, z_{21}, \Phi_2, \Theta_2)$
2. Bulk solvent that is not interfacial or hydrating the proteins will be the same in the bound and unbound ensembles: $C_{bulk} \equiv \int d\boldsymbol{r_{bulk}} \exp(-\beta U(\boldsymbol{r_{bulk}}))$ and $C'_{bulk} \equiv \int d\boldsymbol{r_{bulk,w,m}} \exp\left(-\beta U(\boldsymbol{r_{bulk,w,m}})\right)$ includes membrane solvent.
3. In the unbound ensemble in 2D, the potential energy can be further split into independent contributions from the internal degrees of freedom of each protein and from their position/orientation (height, pitch, roll) in the system: $U(z_1, \Phi_1, \Theta_1, z_{21}, \Phi_2, \Theta_2, \boldsymbol{r_1}, \boldsymbol{r_{s,m,hyd}}, \boldsymbol{r_2}) \cong U(z_1, \Phi_1, \Theta_1) + U(z_{21}, \Phi_2, \Theta_2) + U(\boldsymbol{r_1}, \boldsymbol{r_{s,m,hyd}}, \boldsymbol{r_2})$
4. In the bound ensemble in 2D, the potential energy of protein two relative to protein one can be assumed independent of the position of protein one in the system: $U(z_1, \Phi_1, \Theta_1\, \boldsymbol{r_1}, \boldsymbol{r_{w,m,hyd}}, \boldsymbol{r_2},\ \boldsymbol{R}_{21},\ \boldsymbol{\Omega}_2) \cong U_b(z_1, \Phi_1, \Theta_1) + U(\boldsymbol{r_1}, \boldsymbol{r_{w,m,hyd}}, \boldsymbol{r_2},\ \boldsymbol{R}_{21},\ \boldsymbol{\Omega}_2)$. This assumption is imperfect in 2D, as the potential energy variation with the orientation of protein 2 is likely dependent on protein 1's orientation because the membrane introduces anisotropy in accessible configurations.

$$Z_{21,unb}^{3D} \cong 8\pi^2 (V - V_{cut}) C_{bulk} \int d\boldsymbol{r_1} d\boldsymbol{r}_{hyd} d\boldsymbol{r_2} \exp\left(-\beta U(\boldsymbol{r_1}, \boldsymbol{r_2}, \boldsymbol{r_{hyd}})\right) \qquad (S9a)$$

$$Z_{21,bnd}^{3D} \cong C_{bulk} \int_0^L dz_{21} \int_0^L dy_{21} \int_0^L dx_{21} \int_0^{2\pi} d\Psi_2 \int_0^{2\pi} d\Phi_2 \int_0^{\pi} \sin(\Theta_2)\, d\Theta_2 \int d\boldsymbol{r_1} d\boldsymbol{r}_{hyd} d\boldsymbol{r_2}$$

$$\exp\left(-\beta U\left(\boldsymbol{R}_{21},\ \boldsymbol{\Omega}_2, \boldsymbol{r_1}, \boldsymbol{r_2}, \boldsymbol{r_{hyd}}\right)\right) H(r_{cut} - |\boldsymbol{R}_{21}|). \qquad (S9b)$$

In 2D, using these assumptions, Eq. S6b becomes:

$$Z_{21,unb}^{2D} \cong 2\pi(A - A_{cut}) C'_{bulk} f_{z_1,\Phi_1,\Theta_1} f_{z_{21},\Phi_2,\Theta_2} \int d\boldsymbol{r_1} d\boldsymbol{r_{w,m,hyd}} d\boldsymbol{r_2} \exp\left(-\beta U\left(\boldsymbol{r_1}, \boldsymbol{r_{w,m,hyd}}, \boldsymbol{r_2}\right)\right) \quad (S10a)$$

$$Z_{21,bnd}^{2D} \cong\ C'_{bulk} f_{b,z_1,\Phi_1,\Theta_1} \int_0^{Lmem} dz_{21} \int_0^L dy_{21} \int_0^L dx_{21} \int_0^{2\pi} \mathrm{d}\Psi_2 \int_0^{2\pi} d\Phi_2 \int_0^{\pi} \sin(\Theta_2)\, d\Theta_2$$

$$\int d\boldsymbol{r_A} d\boldsymbol{r_{w,m,hyd}} d\boldsymbol{r_B} \exp\left(-\beta U\left(\boldsymbol{r_1}, \boldsymbol{r_{w,m,hyd}}, \boldsymbol{r_2},\ \boldsymbol{R}_{21},\ \boldsymbol{\Omega}_2\right)\right) H(r_{cut} - |\boldsymbol{R}_{21}|) \qquad (S10b)$$

where

$$f_{z_1,\Phi_1,\Theta_1} = \int_0^{Lmem} \mathrm{d}z_1 \int_0^{2\pi} d\Phi_1 \int_0^{\pi} \sin(\Theta_1)\, d\Theta_1 \exp\left(-\beta U(z_1, \Phi_1, \Theta_1)\right) \qquad (S11)$$

has units of length via the integral over $\mathrm{d}z_1$, and has the same form for protein 2 or the bound state $f_{b,z_1,\Phi_1,\Theta_1}$. This Eq. S11 is the same as the main text Eq. 8. The value of $f$ in 2D will be less than the 3D value due to the membrane restrictions, because the potential energy in 3D is independent of these coordinates and thus integrates to $f_{z_1,\Phi_1,\Theta_1} = 4\pi L$. We note that the 2D $U(z_1, \Phi_1, \Theta_1)$ must therefore measure deviations of the potential energy from the 3D $U(z_1, \Phi_1, \Theta_1) \equiv 0$, which can be achieved by setting its $\min(U) = 0$.

To fully simplify $h$ (Eq. 6 main text) using the partition functions of Eq. S6-S11, we first eliminate the membrane degrees of freedom in 2D by assuming that the interface of the proteins to the membrane contributes the same energetically to the bound and the unbound monomers, $U_{bnd}\left(\boldsymbol{r_{m,hyd}}\right) \cong U_{unb}\left(\boldsymbol{r_{m,hyd}}\right)$. We will also assume the area of volume lost due to the bound state is relatively small compared to the unbound state: $(A - A_{cut})/A \cong 1$ and $V/(V - V_{cut}) \cong 1$ to remove this minor dependence on the system size. With these assumptions and substituting Eq. S6-S11 into Eq. 6 of the main text, we arrive at Eq. 7 of the main text:

$$h \cong \frac{f_{z_1,\Phi_1,\Theta_1} f_{z_{21},\Phi_2,\Theta_2}}{4\pi f_{b,z_1,\Phi_1,\Theta_1}} \frac{\int \exp\left(-\beta U(\boldsymbol{r_1}, \boldsymbol{r_{hyd}}, \boldsymbol{r_2})\right)_{2D,unb} \int \exp\left(-\beta U(\boldsymbol{r_1}, \boldsymbol{r_{hyd}}, \boldsymbol{r_2}, \boldsymbol{R}_{21}, \boldsymbol{\Omega}_2)\right)_{3D,bnd}}{\int \exp\left(-\beta U(\boldsymbol{r_1}, \boldsymbol{r_{hyd}}, \boldsymbol{r_2})\right)_{3D,unb} \int \exp\left(-\beta U(\boldsymbol{r_1}, \boldsymbol{r_{hyd}}, \boldsymbol{r_2}, \boldsymbol{R}_{21}, \boldsymbol{\Omega}_2)\right)_{2D,bnd}} \quad (S12)$$

We compressed the notation of the integrals from Eq. S10 and S11 to highlight that the potential energies now depend on the same variables in 2D and 3D, with the unbound states depending only on the internal coordinates and hydration solvent $(\boldsymbol{r_1}, \boldsymbol{r_{hyd}}, \boldsymbol{r_2})$ while the bound state also depends on the relative position and orientation of protein 2 in the bound state volume ( $\boldsymbol{R}_{21}, \boldsymbol{\Omega}_2$).

**SIII: Constraints to generate the pseudo membrane simulations**

We generated a probability density function $p_{mem}(z_{\text{surf}})$ from the sampled $z_{\text{surf}}$ values from our unbiased trajectories and transform it to a potential of mean force via:

$$U(z_{\text{surf}}) = -k_B T \log\left(p_{mem}(z_{\text{surf}})\right) \quad (S13)$$

The distribution and potential are shown in Fig. S1. The simplest potential is a harmonic potential, and thus we fit our sampled $U(z_{\text{surf}})$ curve to:

$$U(z_{\text{surf}}) = \frac{1}{2} k_z (z_{\text{surf}} - z_0)^2 + cons \quad (S14)$$

Where $k_z$ is the spring constant in units of kJ/mol/nm$^2$ and $z_0$ is the average separation sampled in the simulations. The constant is to constrain the potential to the normalized probability distribution, such that $cons = \frac{k_B T}{2} \ln\left(\frac{2\pi k_B T}{k_z}\right)$. Fitting the simulation data to the harmonic potential produced a best-fit $k_z$ =167 kJ/mol/nm$^2$ and $z_0$ = 1.11 nm for chain A and $k_z$ =160 kJ/mol/nm$^2$ , $z_0$ = 1.32 nm for chain B.

As for the $z_{\text{surf}}$, we can transform the probability distributions we collect for each of the orientational variables $\alpha_{\text{long}}$ and $\alpha_{\text{short}}$ into potentials of mean force. As shown in Fig S1, these potentials are relatively well fit by harmonic potentials of the same form as above. We separately fit the distribution for each monomer, with values collected in Table S2 and Table S3. The spring force constant for the $\alpha_{\text{long}}$ angle is 401 or 550 kJ/mol/rad$^2$ , with the mean value at 1.55 or 1.48 rad for chain A or B, respectively. For the $\alpha_{\text{short}}$ orientation, the force constant was 121 kJ/mol/rad$^2$ or 106 kJ/mol/rad$^2$ , with the mean value being 0.46 or 0.39 rad, for chain A or B, respectively.

**SIV. Metadynamics background and FES calculations.** To briefly explain the parameters defined for a metadynamics simulation, we define the bias potential here. The bias potential is a function of the CVs, $\vec{s} \epsilon \{d_1, d_2\}$ and time $t$, and is expressed as a sum over deposited Gaussian potentials:

$$V(\vec{s}, t) = \sum_{t=\tau_G, 2\tau_G, \ldots}^{t_F} w\, e^{\left(-\frac{(s_1 - d_1(t))^2}{2\sigma_1{}^2}\right)} e^{\left(-\frac{(s_2 - d_2(t))^2}{2\sigma_2{}^2}\right)} \qquad (S15)$$

Here $w$ is the height of the Gaussian potential in kJ/mol, $d_1(t)$ and $d_2(t)$ represent the instantaneous value of the CVs at time t, $\tau_G$is the deposition time of the Gaussian potential, and $\sigma_1$ and $\sigma_2$ are the widths of the Gaussians in the dimension of each CV. After a sufficient long time $t_F$, the free energy $F(\vec{s})$ can be evaluated from the bias potential to with a constant offset:

$$F(\vec{s}) = -V(\vec{s}, t \rightarrow \infty) + C \qquad (S16a)$$

Or in the well-tempered version,

$$F(\vec{s}) = -\frac{\gamma}{\gamma - 1} V(\vec{s}, t \rightarrow \infty) + C_1 \qquad (S16b)$$

where the bias factor $\gamma = \frac{T + \Delta T}{T}$, is the ratio of the effective sampling temperature of the CVs to the system temperature. In our simulations, we estimate the FES from the standard metadynamics using Eq S16a, and the well-tempered part using Eq S16b and then add the corresponding contributions to compute the final FES. The metadynamics are collected in Table S4.

For the solution simulations. The height was initially, $w$=5 kJ/mol, but was reduced to 1 kJ/mol after 28us, to better refine the FES. We switched to well-tempered metadynamics after 106 μs, which allows the height $w$ of the barriers to decrease to converge the FES, $\gamma$ =80. For the membrane simulations, the relaxation time along the CVs is slowed relative to solution due to the reduced diffusion of the proteins on the lipid bilayer. To maintain a relatively frequent deposition time of $\tau_G = 300ps$, we therefore had to reduce the widths of our Gaussians to $\sigma_1 = \sigma_2 = 0.04$nm . Initially we used a height of $w$=5 kJ/mol, but this was reduced to 1 kJ/mol after 96 μs . We switched to well-tempered metadynamics with the bias factor $\gamma = 20$ starting at 156μs . Over the course of the simulation we observed that the well-tempered metadynamics was inhibiting convergence by slowing depositions. We therefore turned off the well-tempered deposition and resumed the standard version with a height of $w$**=**2 kJ/mol after 201 μs. For the pseudo-membrane simulations, the height was initially set to 5kJ/mol, which was then reduced to 1kJ/mol after 12 us. The well-tempered version was switched on after 18 μs with $\gamma = 20$ until 102 μs after which we resumed the standard version with height of $w$**=**1 kJ/mol. These parameters are summarized in Table S4, along with the metadynamics parameters used for the MARTINI 2.0S simulations.

***SV.* Calculating ensemble averages of potential energies*:*** We computed potential energies of both bound and unbound ensembles. To calculate $\langle U \rangle_x$ from an unbiased equilibrium NPT trajectory, we exploit the ergodic principle to take a simple unweighted time average over configurations within each grid point, $U(\vec{s}) = \frac{\sum_{t=1}^{Nt} U(r^N(t))\delta\left(\vec{s}-\vec{s}(r^N(t))\right)}{\sum_{t=1}^{Nt} \delta\left(\vec{s}-\vec{s}(r^N(t))\right)}$, where *Nt* is the total number of configurations from a trajectory. However, to compute ensemble averages from a biased metadynamics trajectory, we must apply a reweighting factor as we average[3]. We use normalized weights,

$$U(\vec{s}) = \frac{\sum_{t=1}^{Nt} U(r^N(t))\delta\left(\vec{s}-\vec{s}(r^N(t))\right)\exp\left(\beta\left[V\left(s(r^N(t)),t\right)-C(t)\right]\right)}{Q_s} \qquad (S17)$$

$$Q_s = \sum_{t=1}^{Nt} \delta\left(\vec{s}-\vec{s}(r^N(t))\right)\exp\left(\beta\left[V\left(s(r^N(t)),t\right)-C(t)\right]\right) \qquad (S18)$$

where $C(t)$ is the time-dependent offset[3] and is calculated using the PLUMED plugin. The potential energy averaged over multiple grid points is given by $\langle U \rangle_x = \frac{\sum_{s_i \in x} U(\vec{s}_i) Q_{s_i}}{\sum_{s_i \in x} Q_{s_i}}$. Normalization reduces numerical issues due to very large values of $V(s(r^N(t)),t) - C(t)$.

For characterizing the system in terms of its deviation from the crystal structure we use two types of RMSD measurements. To evaluate the conformational flexibility within each monomer, we calculate the RMSD value of every frame of the trajectory by aligning each monomer separately to the crystal structure using the MDAnalysis software[4]. This removes the translational and rotational degrees of freedom and then the RMSD is calculated by

$$RMSD\left(m_j\right) = \sqrt{\frac{1}{n}\sum_{i=1}^{n}\left|\boldsymbol{r}_i^{sim,m_j} - \boldsymbol{r}_i^{cry,m_j}\right|^2} \tag{S19}$$

Where $j = 1{,}2$ for each monomer, $n$ is the number of atoms and $\boldsymbol{r}_i^{sim,m_j}, \boldsymbol{r}_i^{cry,m_j}$ are the positions of the corresponding atoms in the simulation and crystal structure respectively. We also calculate $dRMSD$ to assess the dimer configurational states sampled over the course of a simulation. This value is calculated by first aligning monomer 1 to its crystal counterpart. The $dRMSD$ value is then evaluated by calculating the deviation of monomer 2 with respect to its crystal coordinates

$$dRMSD = \sqrt{\frac{1}{n}\sum_{i=1}^{n}\left|\boldsymbol{r}_i^{sim,m_2} - \boldsymbol{r}_i^{cry,m_2}\right|^2} \tag{S20}$$

**Estimates of $\boldsymbol{h_{RIGID}}$ from distributions of height, pitch and roll**

In the Methods we describe how we can numerically calculate the distributions of observed height, pitch, and roll values from our 2D simulations. From these distributions, we construct potential energy functions that are inserted into Eq. 9 (main text) to compute $h_{RIGID}$ from Eq. 10 of the main text. If we assume that each of these potential functions are harmonic in their variable, and we assume the integrals can be extended to $\pm\infty$, these values can be estimated directly from the potential fit parameters:

$$\zeta_1 = \int_0^z e^{-\left(\frac{1}{2k_BT}k_z(z-z_0)^2\right)}dz \;\cong\; \sqrt{\frac{2\pi k_BT}{k_Z}} \tag{S21a}$$

$$f_{\Theta_1} = \int_0^{\pi} \sin(\Theta_1)\, d\Theta_1 \exp\left(-\beta k_\theta \frac{\left(\Theta_1 - \Theta_{1,0}\right)^2}{2}\right) \cong k_B T \frac{}{k_\Theta} \qquad (S21b)$$

$$f_{\Phi_1} = \int_0^{2\pi} e^{-\left(\frac{1}{2k_B T} k_\phi \left(\Phi_1 - \Phi_{1,0}\right)^2\right)} d\Phi_1 \cong \sqrt{\frac{2\pi k_B T}{k_\phi}} \qquad (S21c)$$

These approximations give a sense of how the values vary with the force constants, although we note that the maximal values are $f_{\Theta_1} \leq 2$ and $f_{\Phi_1} \leq 2\pi$, because to compare to 3D we must assume the minimum potential energy is zero. However, we do not report these values in the main text, instead using the numerical integrals over the proper domain range.

### SUPPLEMENTAL FIGURES

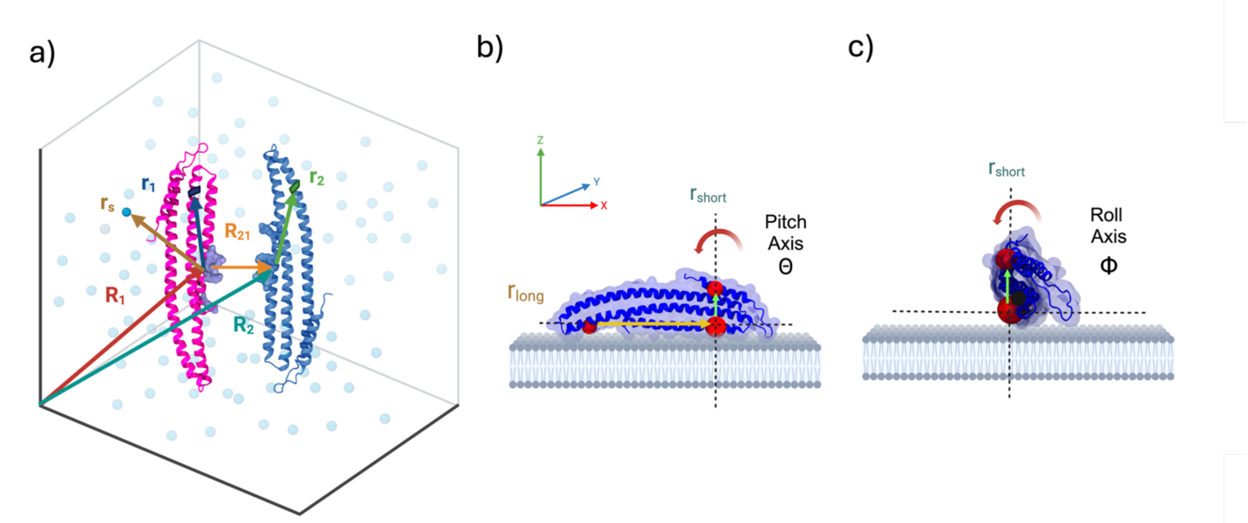


**Figure S1: Definition of the coordinate system and the Tait-Bryan (Euler) angles for rigid-body orientation.** a) The coordinates of the protein rigid body orientation, protein residues and solvent are defined relative to a central point on protein 1, denoted by the vector $R_1$ (orange arrow). The position of protein 2 is relative to protein 1 at $R_{21}$ (yellow arrow). The beads of protein 1 are in the set defined by $\boldsymbol{r_1}^{\boldsymbol{N1}}$ e.g. blue arrow. The beads of protein 2 are in the set defined by $\boldsymbol{r_2}^{\boldsymbol{N2}}$ e.g. green arrow. Solvent beads are in the set defined by $\boldsymbol{r_w}^{\boldsymbol{Nw}}$ (brown arrow). Rigid body orientational angles $\boldsymbol{\Omega}_1$, $\boldsymbol{\Omega}_2$ are defined by the Tait-Bryan/Euler angles of yaw ($\Psi_1$)-not shown, b) the pitch ($\Theta_1$), with the pitch axis coming out of the page and c) the roll ($\Phi_1$), with the roll axis coming out of the page is defined by $r_{long}$. These angles describe intrinsic rotations around the molecule's internal coordinate system $z$, $y'$, $x''$, just as for an airplane (where $y'$ is the axis after the first rotation, $x''$ is after two rotations). Formulas are provided in the Methods for all three of these angles, where yaw measures rotation around the yaw axis $r_{short}$, which is similar to rotation about the global $z$-axis and is thus freely rotating.

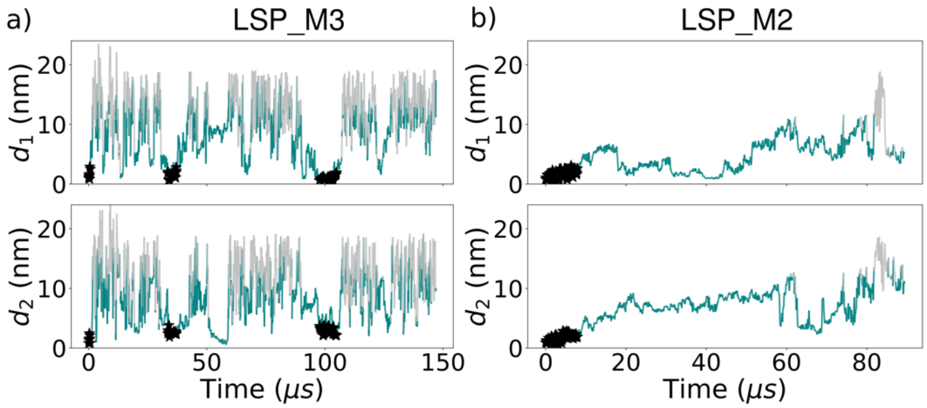


**Figure S2: Collective variables d1 and d2 for the a) LSP_M3.0 and b) LSP_M2.2p pseudo-membrane system versus time.** Gray is the unbound ensemble, green is the bound ensemble, and black stars are the crystal state. Trajectories show many transitions to the bound state, and several to the crystal state (same as Fig 2 for 3D and 2D).

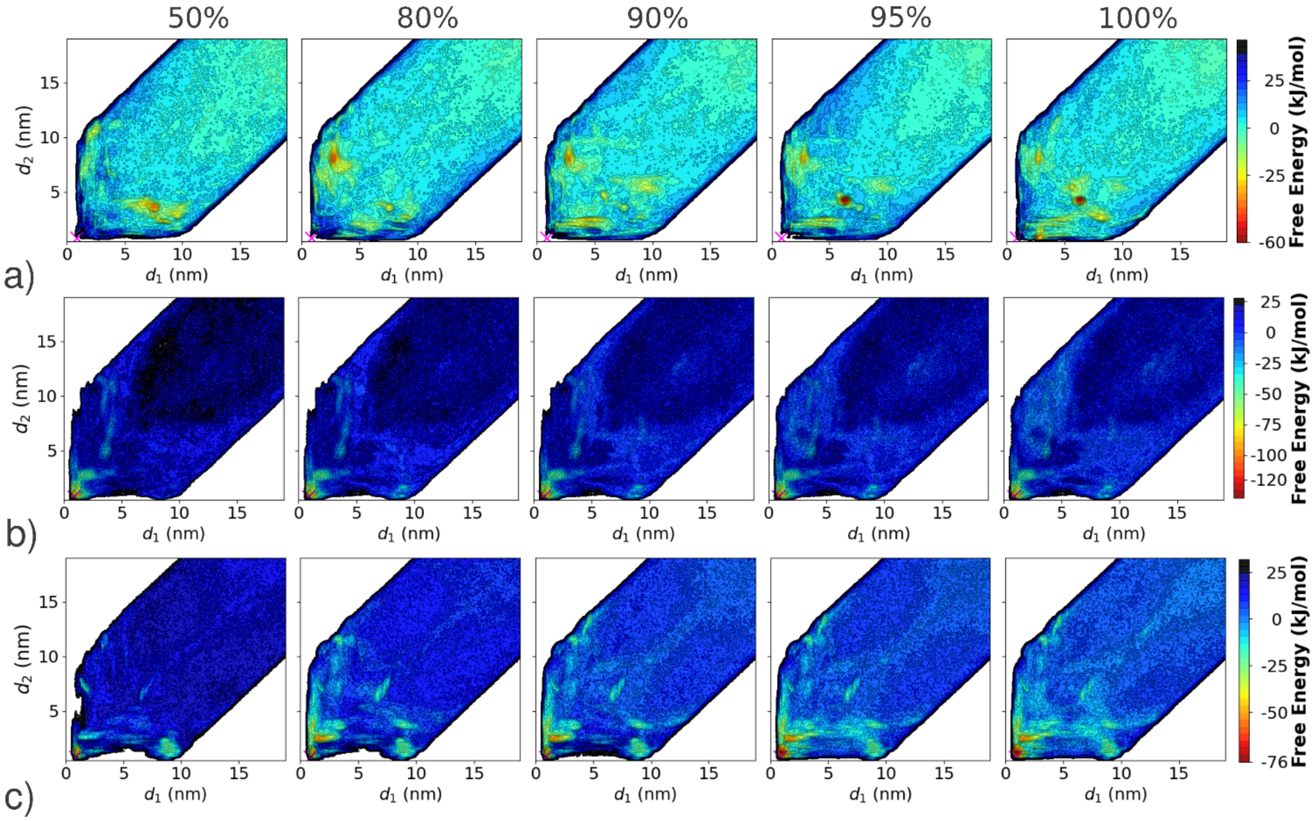


**Figure S3:** Comparing the FES for LSP_M3.0 after 50%, 80%, 90%, 95%, and 100% completeness shows that in both 2D and pseudo-membrane, the crystal state remains the preferred bound configuration, or nearly so. Comparing the surface in 3D at those same points shows that the crystal state remains far from the preferred state, and indeed several nonspecific structures are preferred.

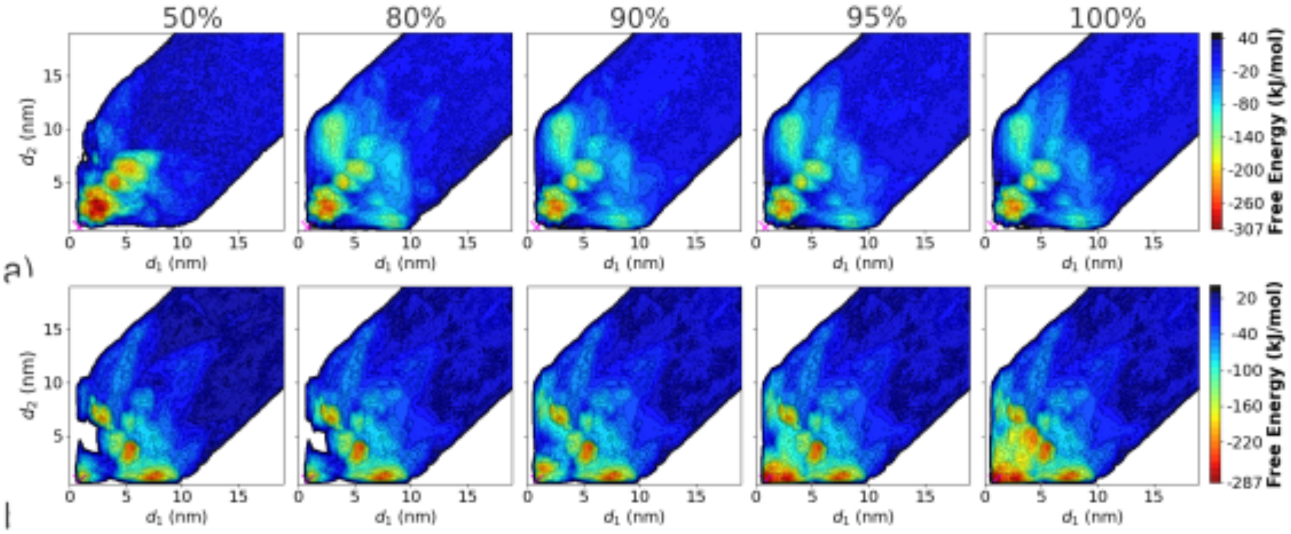


**Figure S4: Comparison of FES progression for LSP_M2.2p.** The FES at different time-points of completion show that the most stable bound structure is closer to the crystal structure (marked by 'X' ) in 2D (lower row) than in 3D (upper row). In both cases, several relatively nonspecific states are sampled, with the 2D states often of comparable free energy as the crystal state despite fluctuations in the free energy around them.

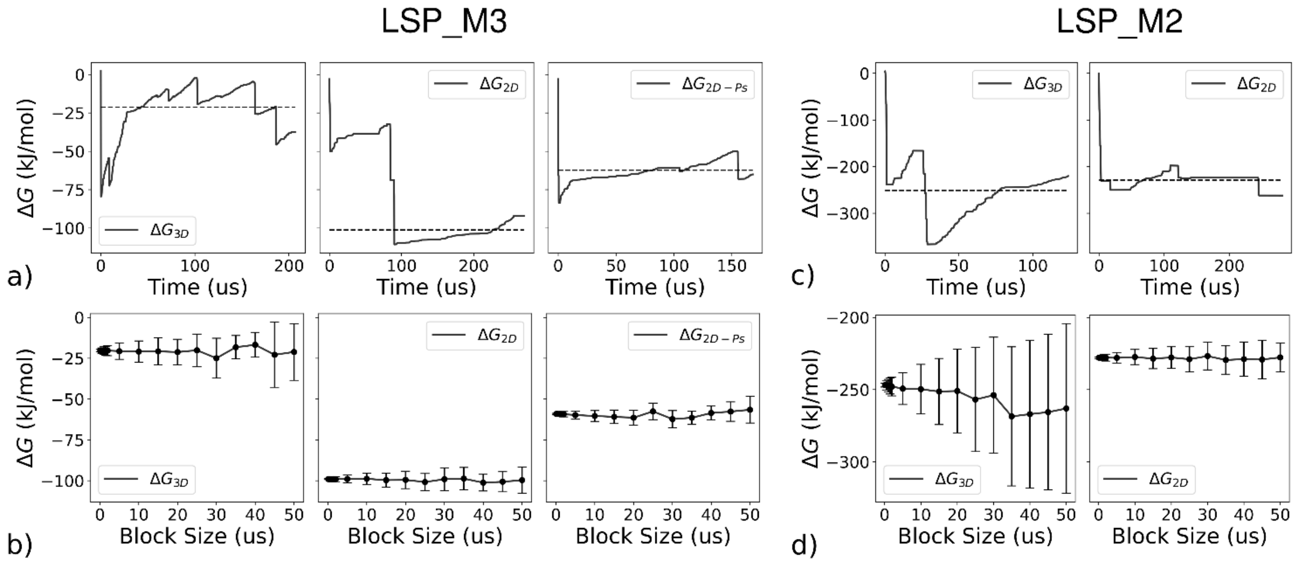


**Figure S5. Convergence and error estimation of *ΔG* from the LSP_M3.0 and LSP_M2.2p free energy surfaces.** a) The time evolution of the free energy difference for the LSP_M3.0 simulations in all three environments. For 3D simulations (left), ΔG shows oscillatory behavior. For 2D simulations however, while there are multiple transitions between the bound and unbound states, the slower diffusion and presence of well-tempered metadynamics make the period of oscillation larger (middle). The 2D-Ps pseudo membrane is on the right. The dashed line represents the averaged ΔG calculated over multiple blocks with block size, 20 μs for 3D and 40 μs for 2D and 2D-Ps. b) The average ΔG and SEM (shown as error bars) obtained for different block sizes over the final 50% of each simulation. c) and d) The free energy differences and averaging data for LSP_M2.2p simulations in 3D (left) and 2D bilayer (right). The 2D pseudo-membrane simulations do not have adequate sampling to assess convergence.

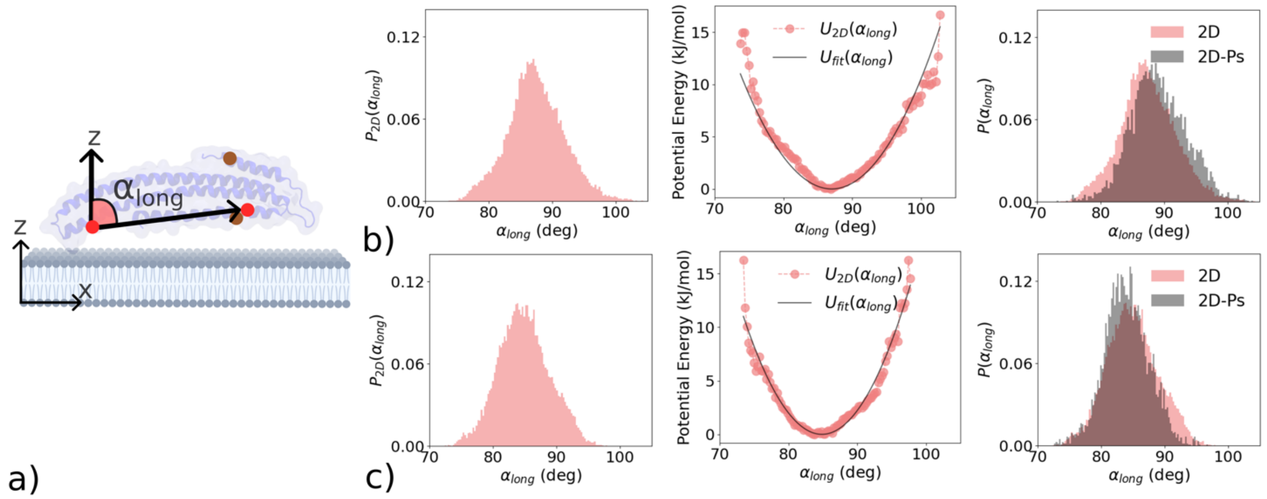


**Figure S6. Calculation of restraint potentials for LSP_M3.0 orientational angle $\alpha_{long}$ a)** Illustration of the beads used to define the vector $r_{long}$ for computing the angle $\alpha_{long}$, which restrains the rotation of the protein picture here around the y-axis, or similar to a pitch angle. A single chain is shown as part of the LSP1 dimer bound to the membrane. b) The first column represents the distribution of $\alpha_{long}$ for a single monomer, from short unbiased simulations of the LSP1 dimer bound to the membrane. The second column represents the fit of the estimated restraint potential to the observed potential energy from the membrane simulations as described in the main Methods via the $ln$ of the probability distribution. The third column compares the distribution of each parameter in the pseudo-membrane simulations with those observed with the actual membrane over a period of ~ 500ns, showing good agreement. c) The process is repeated for the other monomer.

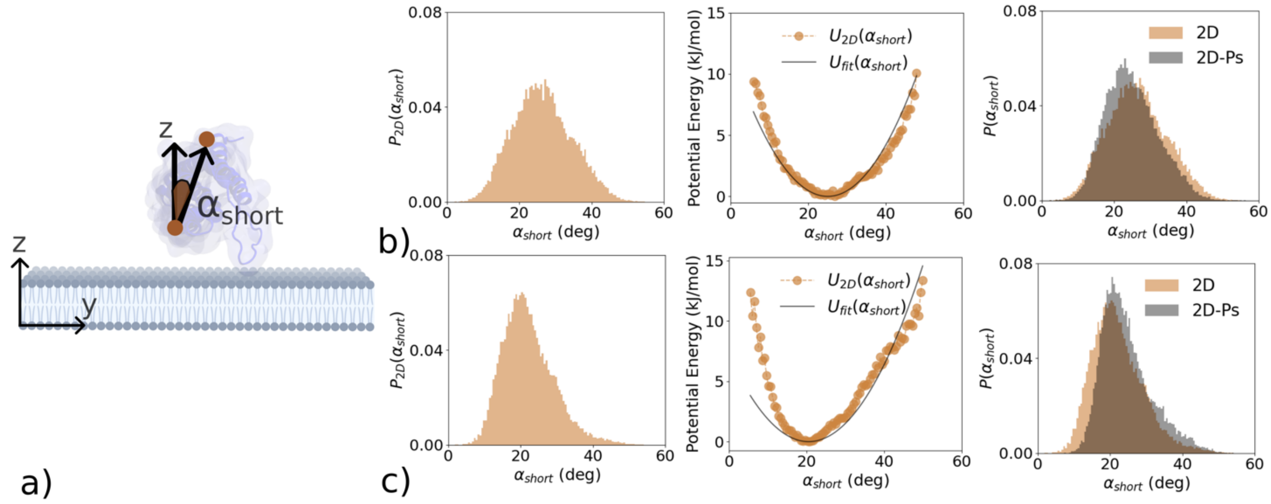


**Figure S7. Calculation of restraint potentials for LSP_M3.0 for orientational angle $\alpha_{short}$. a)** Illustration of the vector used for computing $\alpha_{short}$, which restrains the rotation of the protein around the z-axis defined by the vector $r_{short}$, similar to a roll angle. A single chain is shown as part of the LSP1 dimer bound to the membrane. b) The first column represents the distribution of $\alpha_{short}$ for a single monomer, from short unbiased simulations of the LSP1 dimer bound to the membrane. The second column represents the fit of the estimated restraint potential to the observed potential energy from the membrane simulations (Methods). The third column compares the distribution of each parameter in the pseudo-membrane simulations with those observed with the actual membrane over a period of ~ 500ns showing good agreement. c) Same procedure for the other monomer.

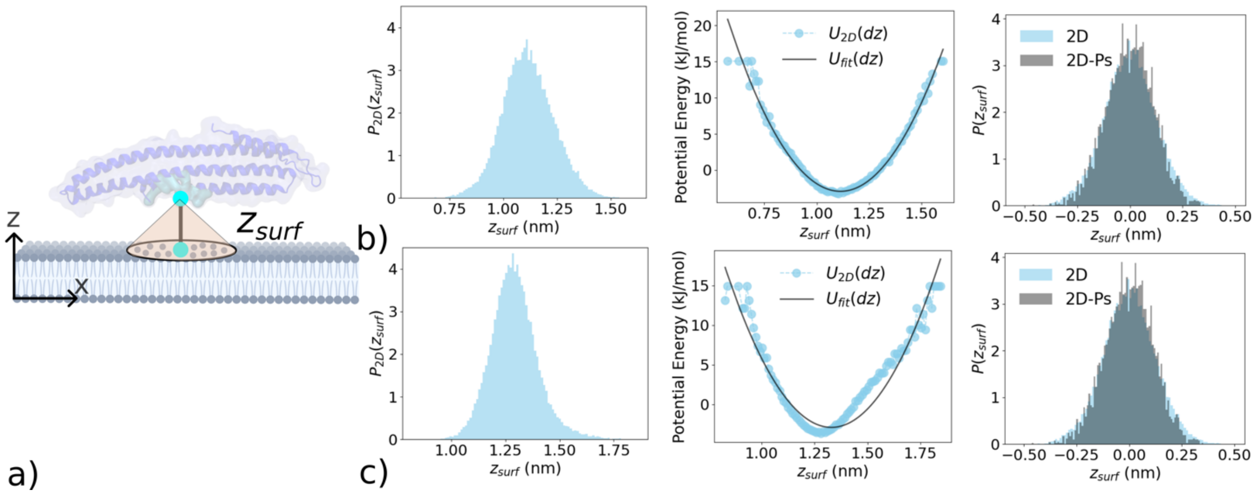


**Figure S8. Calculation of restraint potentials for LSP_M3.0 for the distance $z_{surf}$. a)** Illustration of the method used for computing $z_{surf}$, which restrains the displacement of the protein from the membrane in z-direction. The two beads represent the COM of membrane binding residues on the protein (top) and the COM of phospholipid headgroups which are within 1.2 nm from the membrane binding residues. b) The first column represents the distribution of the $z_{surf}$, for a single monomer, from short unbiased simulations of the LSP1 dimer bound to the membrane. The second column represents the fit of the estimated restraint potential to the observed potential energy from the membrane simulations (Methods). The third column compares the distribution of each parameter in the pseudo-membrane simulations with those observed with the actual membrane over a period of ~ 500ns, showing good agreement. c) Same procedure for the other monomer.

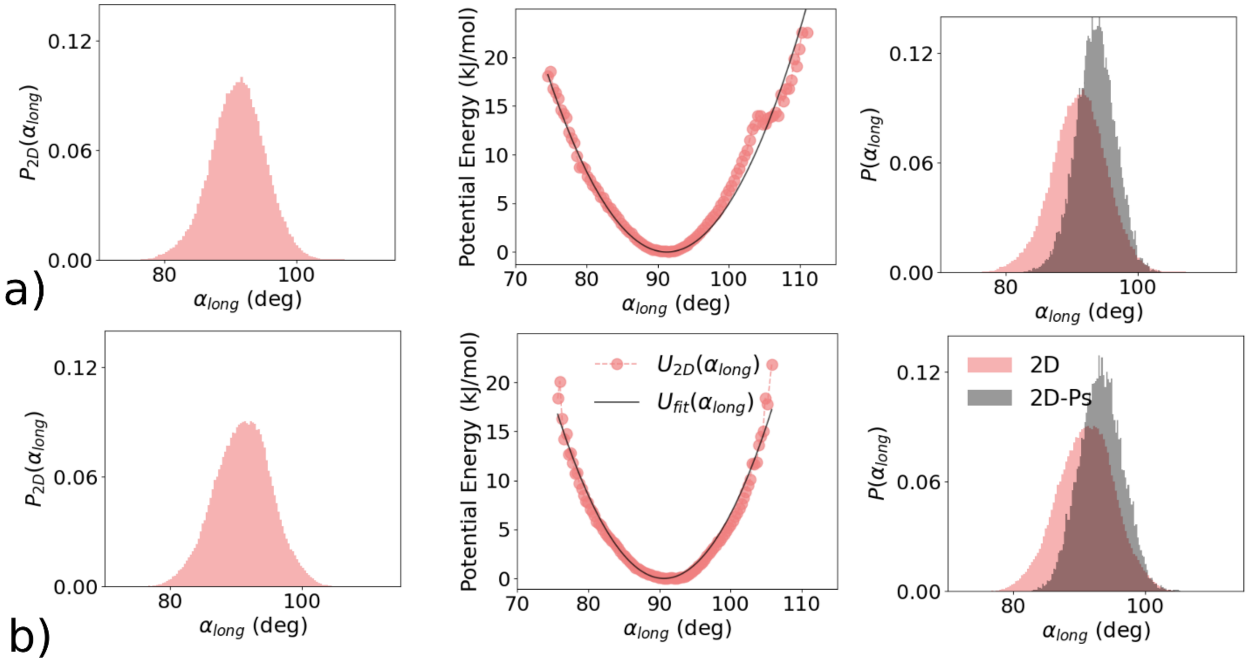


**Figure S9. Calculation of restraint potentials for LSP_M2.2p for orientational angle $\boldsymbol{\alpha_{long}}$ a)** Calculating restraint potential for monomer A - Membrane distributions of $\alpha_{long}$ (left), are converted to a potential energy function (middle) and fitted to a harmonic potential. The resultant distributions are plotted with the membrane distributions (right). **b)** Restraint potential calculations for monomer B.

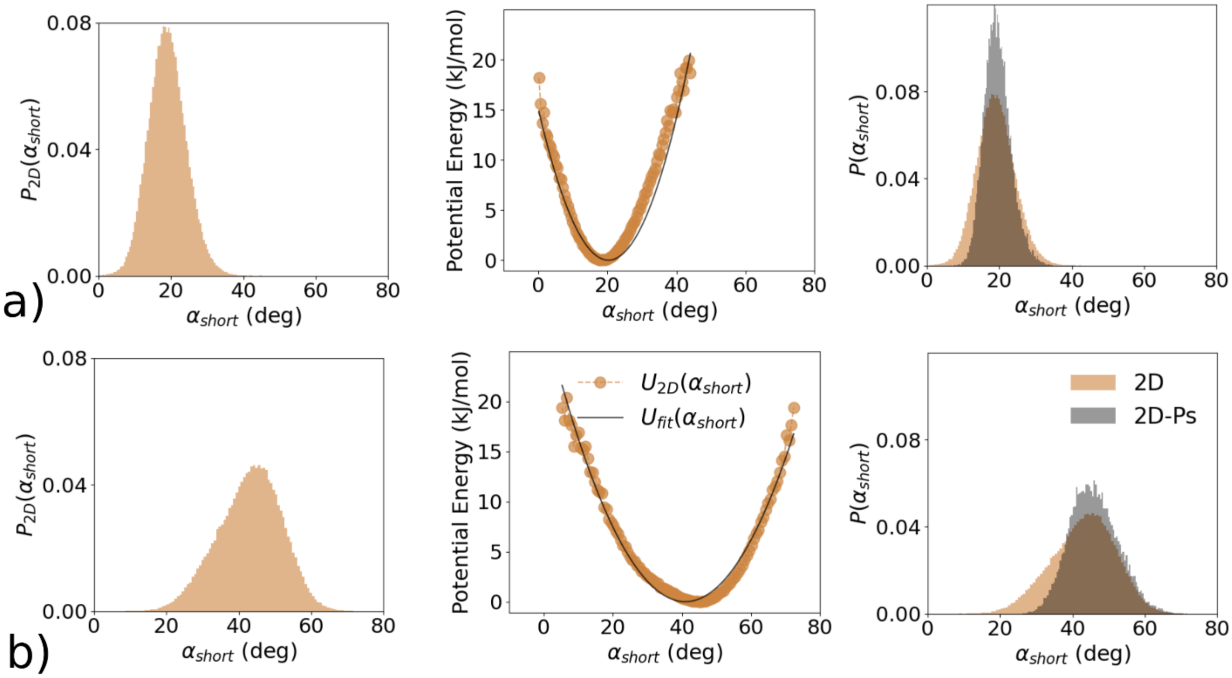


**Figure S10. Calculation of restraint potentials for LSP_M2.2p for orientational angle $\boldsymbol{\alpha_{short}}$ a)** Calculating restraint potential for monomer A - Membrane distributions of $\alpha_{short}$ (left), are converted to a potential energy function (middle) and fitted to a harmonic potential. The resultant distributions are plotted with the membrane distributions (right). **b)** Restraint potential calculations for monomer B.

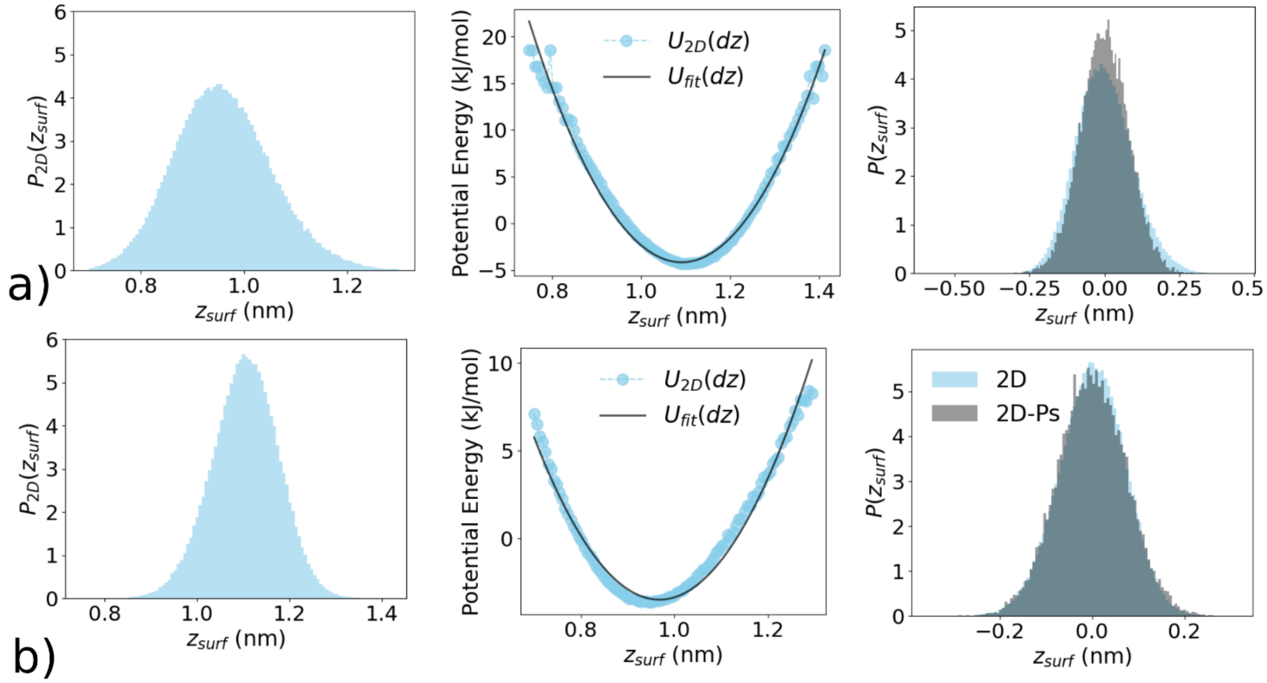


**Figure S11 Calculation of restraint potentials for LSP_M2.2p for the distance $z_{surf}$ a)** Calculating restraint potential for monomer A - Membrane distributions of $z_{surf}$ (left), are converted to a potential energy function (middle) and fitted to a harmonic potential. The resultant distributions are plotted with the membrane distributions (right). **b)** Restraint potential calculations for monomer B.

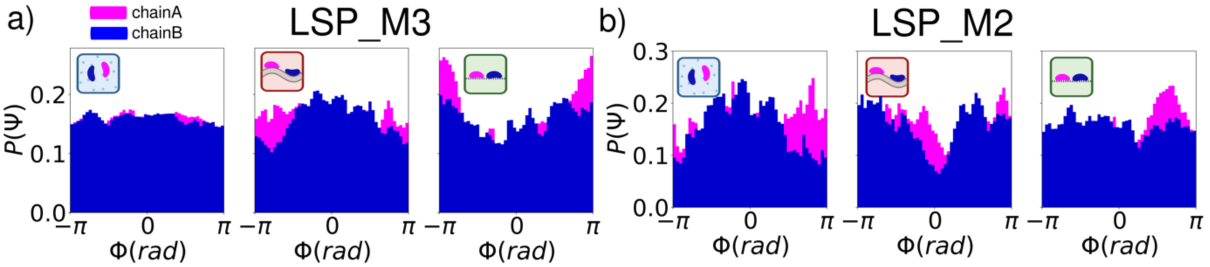


**Figure S12. Distribution of the Yaw euler angles.** Yaw angles which represent rotation about the z-axis, shows unrestricted sampling for all 3 systems in a) LSP_M3.0 and b) LSP_M2.2p.

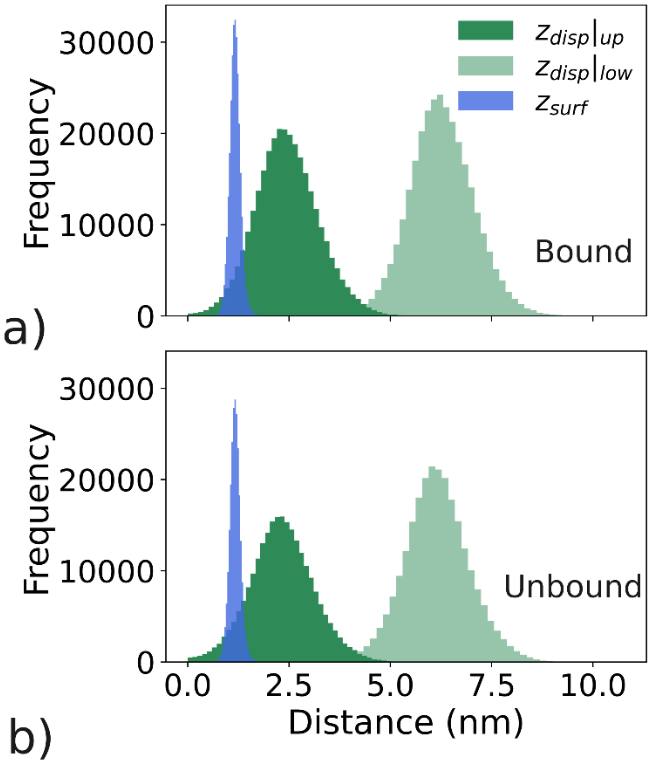


**Figure S13. For LSP_M3.0, we compare how the fluctuations in the z-dimension depend on the choice of variables.** For the $z_{surf}$ metric used to parameterize the pseudo membrane constraint, the separation reports a minimum distance between protein and the membrane surface and is more tightly constrained (blue). When we instead measure the distance in z between the protein zCOM and the z position of the membrane at the same x-y coordinates, the distribution is much broader (green curves), better capturing the fluctuations of the protein and membrane system collectively. The distances to the upper leaflet (dark green) are similar in width to the distances to the lower leaflet (light green) but slightly broader.

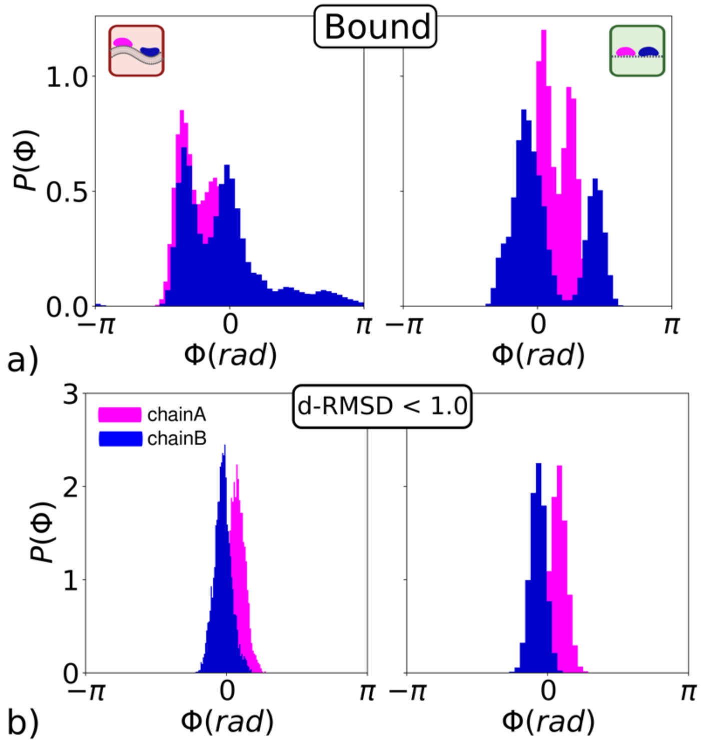


**Fig S14. Distribution of Roll angle for bound ensemble and native dimer specific configurations for LSP_M2.2p.** The formation of stable non-specific interfaces in LSP_M2.2p biases the rotational degree of freedom as evaluated by the roll angle. a) Distribution of the roll angles for all configurations belonging to the bound ensemble (similar to Fig 3g for 2D-Ps, Main text). The distributions stray significantly between the 2D and 2D-Ps simulations and are significantly skewed. b) Here we show the same distribution but for only a subset of configurations closest to the native LSP_M2.2p state with dRMSD < 1.0 nm. This shows that a narrower distribution of roll angles are reproduced for the native dimer state, more similar to what is seen in LSP_M3.0 (Fig 3 Main text).

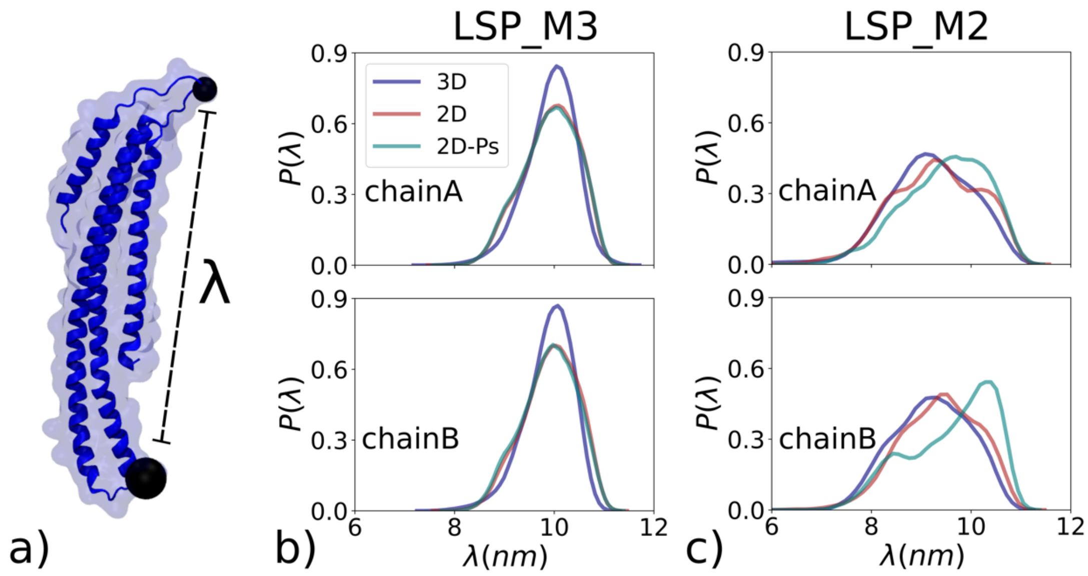


**Figure S15. End to End distance measurements for monomers in LSP_M3.0 and LSP_M2.2p**. a) We calculate the end-to-end distance, defined as the distance between the beads 84THR and 159PRO, for b) LSP_M3.0 c) LSP_M2.2p simulations. The LSP_M2.2p proteins show a significant sampling of conformations with smaller $\lambda$ ($< 8.0\ nm$), which indicates that both chains undergo bending not observed for LSP_M3.0. These contrasting conformations are driven by strong protein-protein contacts formed between the LSP_M2.2p dimers.

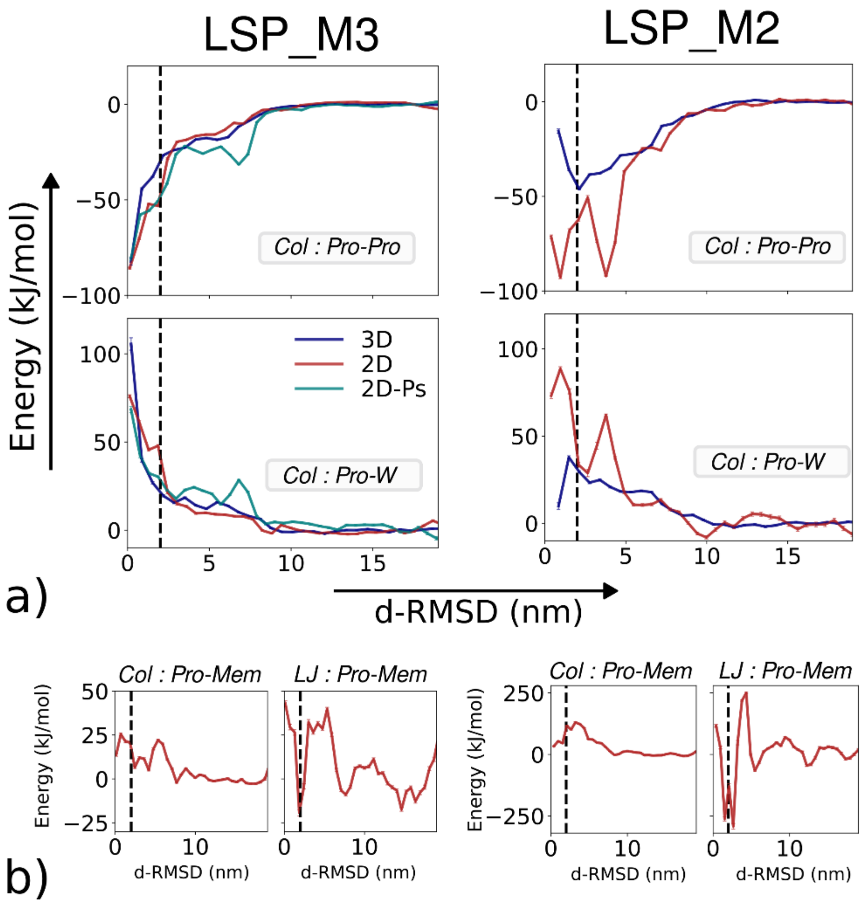


**Figure S16. Potential Energy terms for distinct environments as they depend on proximity to the native dimer state (d-RMSD).** a) The Columbic interaction energy between Protein-Protein interface (top) shows favorable energies for bound LSP1 dimers while Protein-Water interface interactions become unfavorable. While all 3 environments in MARTINI 3 have similar energetic contributions to the native state, the 3D MARTINI 2 simulations display a significant destabilization of the native state as compared to 2D MARTINI 2. b) The non-bonded interactions between protein and membrane do not show significant contribution in terms of kJ/mol when compared to the non-bonded protein-water interactions. However, the LJ interactions indicate that the membrane environment can favor transition to the native state.

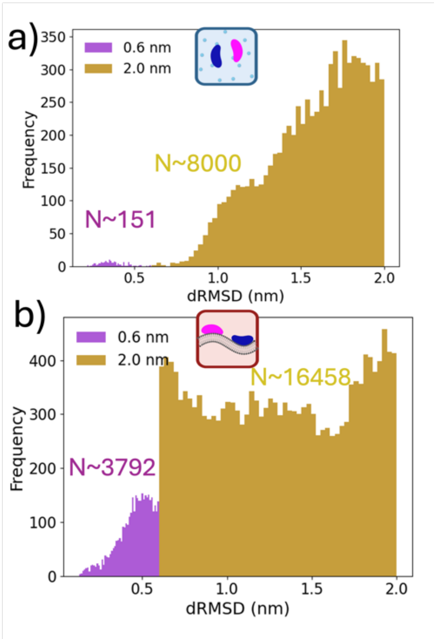


**Figure S17. Distribution of sample dRMSD values for the configurations in near-native ensembles for LSP_M3.0 in 3D versus 2D. a)** Solution configurations. Observed frequencies are out of N=330,000 configurations from the full bound ensemble. For the two classes, with dRMSD < 0.6 nm (purple) and 0.6< dRMSD < 2.0 nm (gold), we see that the LSP_M3.0 distribution is biased towards structures with larger dRMSD values. **b)** In contrast, for the 2D bilayer systems, far more structures have dRMSD< 0.6nm, and the 2.0nm class is much more uniformly distributed in dRMSD values. This is despite having a similar number of N=368,000 configurations from the full bound ensemble. The 2D-Ps values were sampled from N=450,000 configurations, with 816 configurations found in the 0.6nm class (in between 3D and 2D), and 19,494 configurations in the 2.0nm class (more than both 3D and 2D).

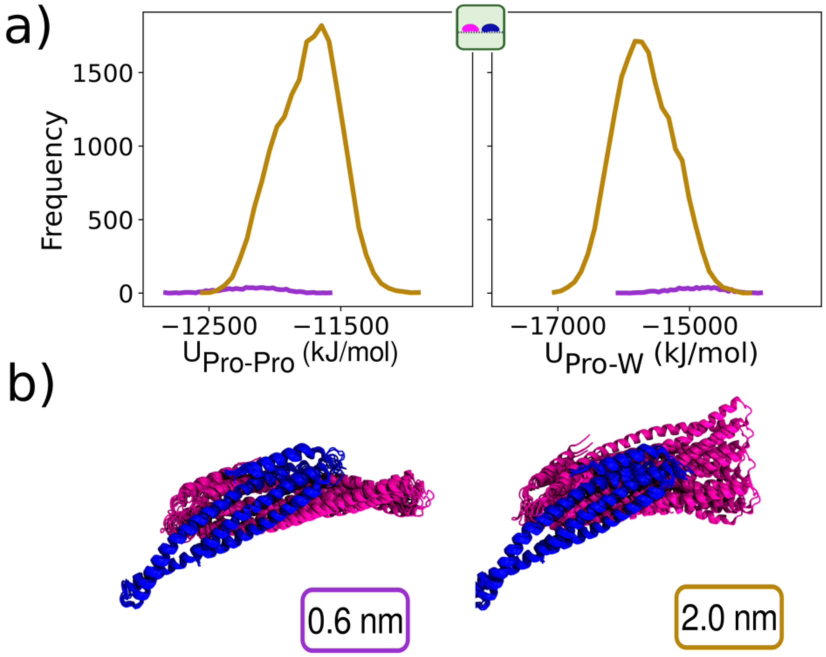

**Figure S18. Energetic and configurational sampling of the native dimer state in 2D-Ps for LSP_M3.0.** a) The Lennard Jones energy distributions for the Protein-Protein and Protein-Water interactions are shown for two mutually exclusive sets of configurational states with dRMSD < 0.6nm (purple) and 0.6nm <dRMSD <2.0 nm (gold). b) The configurations for the pseudo-membrane are similar to those observed on the membrane surface.

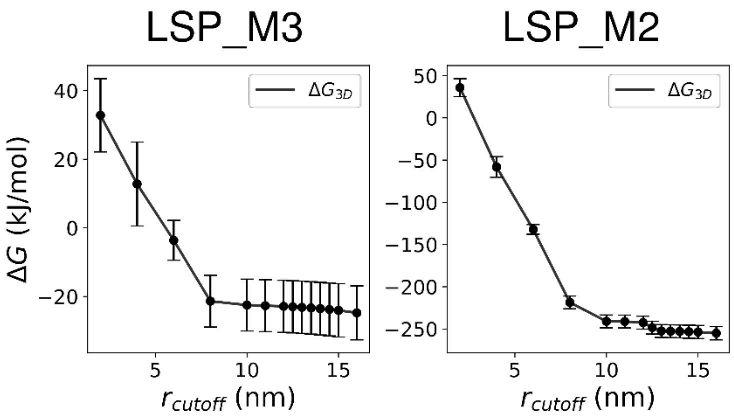


**Figure S19. ΔG dependence on cutoff distance for unbound states.** The ΔG for each system is calculated as the ratio of partition functions between the bound and the unbound state where the bound state is defined as $d_1 < r_{cutoff}$ and $d_2 < r_{cutoff}$ and all other configurations are classified as the unbound state.

### SUPPLEMENTAL TABLES

| **Table S1: Lengthscale $h$ calculations and numerical integration of Eq 9 (Main text)** | | | | | | |
|---|---|---|---|---|---|---|
| **System** | $\zeta_{21}$ | $f_{\Theta 2}$ | $f_{\Phi 2}$ | $h_{RIGID}$ | $h$ (from $\langle \Delta G \rangle$) | $K_D^{2D}$ |
| 2D: LSP_M3.0 | 1.81 nm | 0.243 rad | 0.905 rad | 3 x$10^{-2}$ nm | ~1x$10^{-12}$ nm | $2x10^{-20}$/nm$^2$ |
| 2D-Ps: LPS_M3.0 | 0.413 nm | 0.161 rad | 0.876 rad | 4 x$10^{-3}$ nm | 4.1 x$10^{-6}$ nm | $3x10^{-14}$/nm$^2$ |
| 2D:LPS_M 2.2p | 1.611 nm | 0.215 rad | 0.649 rad | 1.8x$10^{-2}$nm | 1 x$10^{5}$ nm | $3x10^{-42}$/nm$^2$ |

| Table S2: Parameters for the three restraint potentials in pseudo-membrane simulations for LSP_M3.0 (MARTINI 3.0) | | | | |
|---|---|---|---|---|
| | **Chain A** | **95% CI** | **Chain B** | **95% CI** |
| $k_z\ (\frac{kJ}{mol\ nm^2})$ | 167 | (157,165) | 160 | (152,168) |
| $z_{surf}\ (nm)$ | 1.11 | (1.109,1.117) | 1.32 | (1.310,1.329) |
| $k_\theta\ (\frac{kJ}{mol\ rad^2})$ | 401 | (378,414) | 550 | (532,567) |
| $\theta_0\ (rad)$ | 1.55 | (1.545, 1.554) | 1.48 | (1.478, 1.483) |
| $k_\phi\ (\frac{kJ}{mol\ rad^2})$ | 121 | (116,128) | 106 | (88,123) |
| $\phi_0\ (rad)$ | 0.46 | (0.452, 0.467) | 0.39 | (0.360, 0.420) |

| Table S3: Parameters for the three restraint potentials in pseudo-membrane simulations for LSP_M2.2p (MARTINI 2)s | | | | |
|---|---|---|---|---|
| | **Chain A** | **95% CI** | **Chain B** | **95% CI** |
| $k_z\ (\frac{kJ}{mol\ nm^2})$ | 334 | (327,341) | 397 | (389,405) |
| $z_{surf}\ (nm)$ | 1.0 | (0.997,1.002) | 1.09 | (1.088,1.092) |
| $k_\theta\ (\frac{kJ}{mol\ rad^2})$ | 428 | (417,438) | 492 | (480,505) |
| $\theta_0\ (rad)$ | 1.59 | (1.589, 1.595) | 1.58 | (1.579, 1.584) |
| $k_\phi\ (\frac{kJ}{mol\ rad^2})$ | 240 | (230,250) | 111 | (109,113) |
| $\phi_0\ (rad)$ | 0.35 | (0.0.346, 0.0.361) | 0.71 | (0.704, 00.718) |

| **Table S4: Metadynamics parameters and schedule for all simulations** | | | | | | | |
|---|---|---|---|---|---|---|---|
| **Force-field** | **Environment** | **Time span** (μs) | **Std/WT MetaD** | $\sigma_1$, $\sigma_2$ $(nm)$ | $\tau_G$ (ps) | **Total time** (μs) | **Box size** (nm) |
| M3 | Solution/3D | 0 – 28<br>28 – 100<br>100 – 208 | Std ($w$ = 5kJ/mol)<br>Std ( $w$ = 1 kJ/mol)<br>WT ( $b.f.$ = 80) | 0.1 | 280 | 208 | 37 x 37 x 37 |
| M3 | Membrane/2D | 0 – 96<br>96 – 159<br>159 – 216<br>216 – 249<br>249 – 267 | Std ( $w$ = 5 kJ/mol)<br>Std ( $w$ = 1 kJ/mol)<br>WT ( $b.f.$ = 20)<br>WT ( $b.f.$ = 100)<br>Std ( $w$ = 2.5 kJ/mol) | 0.03 | 300 | 267 | 37 x 37 x 18 |
| M3 | Pseudo-Membrane/2D | 0 – 12<br>12 - 18<br>18 – 102<br>102 – 186 | Std ( $w$ = 5 kJ/mol)<br>Std ( $w$ = 1kJ/mol)<br>WT ( $b.f.$ = 10)<br>Std ( $w$ = 1 kJ/mol) | 0.03 | 300 | 186 | 37 x 37 x 37 |
| M2 | Solution/3D | 0 – 61<br>61 – 123 | Std ( $w$ = 10 kJ/mol)<br>Std ( $w$ = 3 kJ/mol) | 0.1 | 500 | 123 | 38 x 38 x 38 |
| M2 | Membrane/2D | 0 – 151<br>151 – 157<br>157 – 238<br>238 – 286 | Std ( $w$ = 5kJ/mol)<br>Std ( $w$ = 1kJ/mol)<br>WT (bf = 20)<br>Std ( $w$ = 2 kJ/mol) | 0.04 | 300 | 286 | 37 x 37 x 18 |
| M2 | Pseudo-Membrane/2D | 0 – 12<br>12 – 18<br>18 – 57<br>57 – 87 | STD ( $w$ = 5 kJ/mol)<br>WT ( $b.f.$ = 20)<br>Std ( $w$ = 1 kJ/mol)<br>Std ( $w$ = 2.5 kJ/mol) | 0.04 | 300 | 87 | 37 x 37 x 18 |
